# Fast 3D Clear: A Fast, Aqueous, Reversible Three-Day Tissue Clearing Method for Adult and Embryonic Mouse Brain and Whole Body

**DOI:** 10.1101/2021.06.25.449994

**Authors:** Stylianos Kosmidis, Adrian Negrean, Alex Dranovsky, Attila Losonczy, Eric R. Kandel

## Abstract

To date, a variety of optical clearing methods have emerged that serve as powerful tools to study intact organs and neuronal circuits. Here we describe a newly developed, aqueous clearing protocol named “Fast 3D Clear”, which relies on tetrahydrofuran (THF) for tissue delipidation, and iohexol (Histodenz) for clearing, such that tissues can be imaged under immersion oil in light sheet imaging systems. Fast 3D Clear requires three days to achieve high transparency of adult and embryonic mouse tissues, while maintaining their anatomical integrity, and preserving a vast array of transgenic and viral/dye fluorophores, such as GCaMP3/6, tdTomato, Fast Blue, and IRF670. A unique advantage of Fast 3D Clear is its complete reversibility and thus compatibility with tissue sectioning and immunohistochemistry. Fast 3D Clear can be easily and quickly applied to a wide range of biomedical studies, thereby greatly facilitating the acquisition of high-resolution, two - and three -dimensional images.

## Introduction

Since tissue clearing was first described by Werner Spalteholz over a century ago (1914) (Spalteholz, 1914; Steinke and Wolff, 2001), several optical clearing techniques have been introduced that eliminate labor-intensive histological sectioning and facilitate studies on neuronal development, morphology, and connectivity. These techniques include, but are not limited to 3DISCO (Erturk et al., 2012), iDISCO (Renier et al., 2014), uDISCO (Pan et al., 2016), FDISCO (Qi et al., 2019), FluoClearBABB (Schwarz et al., 2015), PEGASOS (Jing et al., 2018), CLARITY (Chung and Deisseroth, 2013) PACT-PARS (Yang et al., 2014), CUBIC (Tainaka et al., 2014), BONE Clear (Wang et al., 2019) and SWITCH (Murray et al., 2015).

Clearing methods can be categorized as organic solvent-based (i.e 3DISCO, iDISCO, uDISCO, FDISCO, FluoClearBABB, PEGASOS), or as aqueous (i.e CLARITY, PACT-PARS, CUBIC), each one having its advantages and limitations. Organic solvent-based protocols provide high-level tissue transparency in only three to four days, with the exception of FluoClearBABB that requires ten days. The main disadvantages of these protocols include bleaching of fluorescent protein labels (3DISCO), long antibody incubation times (iDISCO), complexity in their operation (uDISCO), the toxicity of some organic solvents, and tissue shrinkage that can impede high-resolution imaging (FDISCO). On the other hand, aqueous methods are simple in their application, and can preserve fluorescent proteins. However, these protocols often require specific equipment (CLARITY), and the clearing process is slow, taking up to fourteen (CUBIC) or seventeen days (PACT-PARS).

We have built upon the powerful techniques described previously to develop a new method of whole tissue clearing that integrates the desirable simplicity, speed, versatility, and high-transparency, without sacrificing safety, tissue integrity, and the preservation of endogenous fluorescence. The proposed Fast 3D Clear results in highly transparent tissues such as whole adult mouse bodies and brains, embryonic spinal cord, and retina withing three days, requiring only four solutions and seven steps. The refractive-index matching aqueous clearing and imaging solution formulation does not produce toxic vapors and is compatible with standard microscopy instrumentation and optics. The tissue morphology and size are not compromised, while endogenous fluorescent labels with emission spanning from blue to far red are preserved for more than seven months. Importantly, the clearing procedure of Fast 3D Clear is reversible, as tissues can be returned to their previous non-transparent state and are suitable for further processing with immunohistochemistry/immunofluorescence. Using Fast 3D Clear together with light sheet or confocal microscopy, we were able to observe neurons of the adult mouse hippocampus and map dentate gyrus inputs from the entorhinal and perirhinal cortex in virally-expressed fluorescent protein labeled brains.

## Results

### Fast 3D Clear achieves high tissue transparency in brains and whole adult mice and embryos

Fast 3D Clear consists of a minimal number of steps and requires only three days to achieve complete transparency (Figure 1A). We chose Tetrahydrofuran (THF) as a dehydration/ de-lipidation agent to develop a fast and reversible clearing method. THF has been previously used in whole tissue clearing (Erturk et al., 2012) and it can rapidly infiltrate and conserve soft tissues for immunohistochemistry (Haust, 1959). To avoid bleaching of genetically expressed reporters such as GFP and tdTomato, we used THF at pH 9-9.5, which has been shown to reduce fluorescence quenching (Qi et al., 2019). To further avoid deterioration of fluorescence and overall tissue integrity (i.e., shrinkage), as happens with the use of organic solutions (100% THF), we reversed THF-induced dehydration. We achieved this by gradually decreasing THF concentrations to water, leading to complete restoration of tissue size (Figure 1B,1C). We noticed that prolonged (overnight) incubation/washing of the brain with deionized water after THF treatment causes a ∼25% linear expansion of the tissue compared to its original size (Supplemental Figure 1A). To maintain this tissue expansion along with the fluorescence, we incorporated urea into the iohexol (Histodenz)-based clearing solution (One-way ANOVA F (2.24) = 44.11 p<0.0001) (Fixed: 1.532 cm, ±0.07, Cleared no urea 1.593 cm, ±0.07 Fixed urea:1.890cm ±0.1056 SEM) (Figure 1D and Supplemental Figure 1C, 1D). We used this aqueous clearing solution to preserve and visualize the cleared tissue in a refractive index matched non-toxic Cargille Type A immersion oil with RI = 1.515 (Supplemental Figure 1B). Fast 3D Clear resulted in intact adult mouse brains with high transparency (Figure 1C, 1D) compared to fixed brains (Figure 1B and Supplemental Figure 1B). We next tested the Fast 3D Clear method in whole adult mice and mouse embryos. Fast 3D Clear was able to produce sufficiently transparent E18.5 mouse embryos and P24, 1-month, 3-month adult whole mice (Figure 1E-1J), as well as whole soft organs including liver, kidneys, lungs, and intestine (Supplemental Figure 1E-1H) while maintaining fluorescence in the peripheral organs without affecting the background. (Supplemental Figure 1I-1M). Thus, Fast 3D Clear is a simple procedure that leads to high transparency in a wide variety of tissues, including the central nervous system.

**Figure 1.**
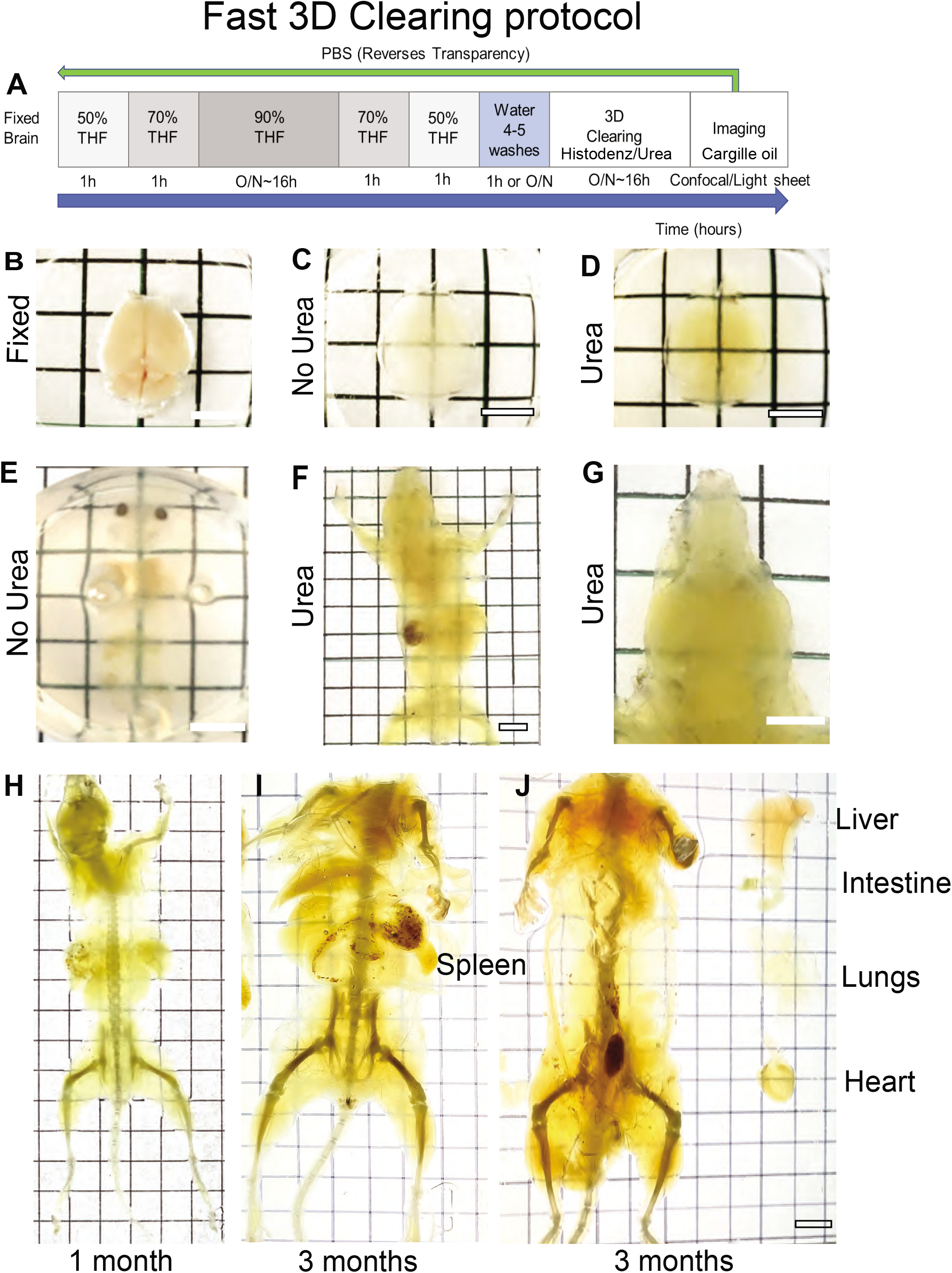
Overview of Fast 3D Clear. A) Schematic of the Fast 3D Clear procedure. B-D) Transparency of adult mouse brains in immersion oil with refractive index RI = 1.515 at 589.3 nm (sodium D-line). B) Fixed brain, C) Mouse brain cleared without urea addition, D) Mouse brain cleared with urea. E) E18.5 mouse embryo without urea. F) Whole adult (P24) mouse cleared with urea, and G) Dorsal view of the head. H) Whole adult 1-month-old mouse cleared with urea. I-J) Whole adult 3-month-old mouse cleared with urea with (I) or without (J) the internal organs. (Scale bar 6 mm).

### Comparison of Fast 3D Clear with other clearing methods

To assess our method’s efficiency and simplicity, we subjected brain hemispheres derived from the same animal to Fast 3D Clear and FDISCO, or Fast 3D Clear and RTF (Figure 2A, 2B and Supplemental Figure 2A). We observed superior clearing of Fast 3D Clear compared to RTF (Supplemental Figure 2A) and similar transparency with the FDISCO method (Figure 2A, 2B). Importantly, there was no noticeable difference in the visual transparency of the cleared tissue when transferred from the aqueous clearing solution to the Cargille immersion oil with RI of 1.515 (Figure 2C, 2E). However, there were considerable differences in the size of tissue cleared with Fast 3D Clear versus FDISCO (Figure 2A-2C). While FDISCO shrinks the tissue, Fast 3D Clear may lead to tissue expansion (Supplemental Figure 2B) and therefore may provide higher magnification of the region of interest. We also used Wintergreen oil as an imaging medium with a higher RI (RI-1.536) (Figure 2D). Although Wintergreen oil led to tissue transparency similar to Cargille oil, its use disintegrated plastic holders. Nevertheless, our results highlight the potential compatibility of Fast 3D Clear with other commercially available immersion oils with higher RI. We next tested whether there were differences between Fast 3D Clear and FDISCO in fluorescence preservation and background fluorescence, using brains from GCaMP3-CaMK2-Cre transgenic animals (Tsien et al., 1996; Zariwala et al., 2012). Using confocal (Figure 2F, 2H and Supplemental Figure 2D, 2E) and light sheet microscopy (Figure 2G, 2I), we measured the signal-to-noise ratio (SNR) using 525/50 nm laser excitation. Fast 3D Cleared hemispheres had a higher SNR (Figure 2J and Supplemental Figure 2C) than the respective FDISCO counterparts, Importantly, Fast 3D Clear was able to preserve fluorescence signal, similar to the one obtained with traditional immunohistochemistry (Supplemental Figure 2D-2I), and the GCaMP3 signal was preserved for at least seven months in Cargille immersion oil (Supplemental Figure 2J-2L). Overall, the application of Fast 3D Clear resulted in equal or better tissue transparency and SNR compared to FDISCO and RTF, and maintenance of the fluorescent signal similar to that of samples processed with classical immunohistochemistry.

**Figure 2.**
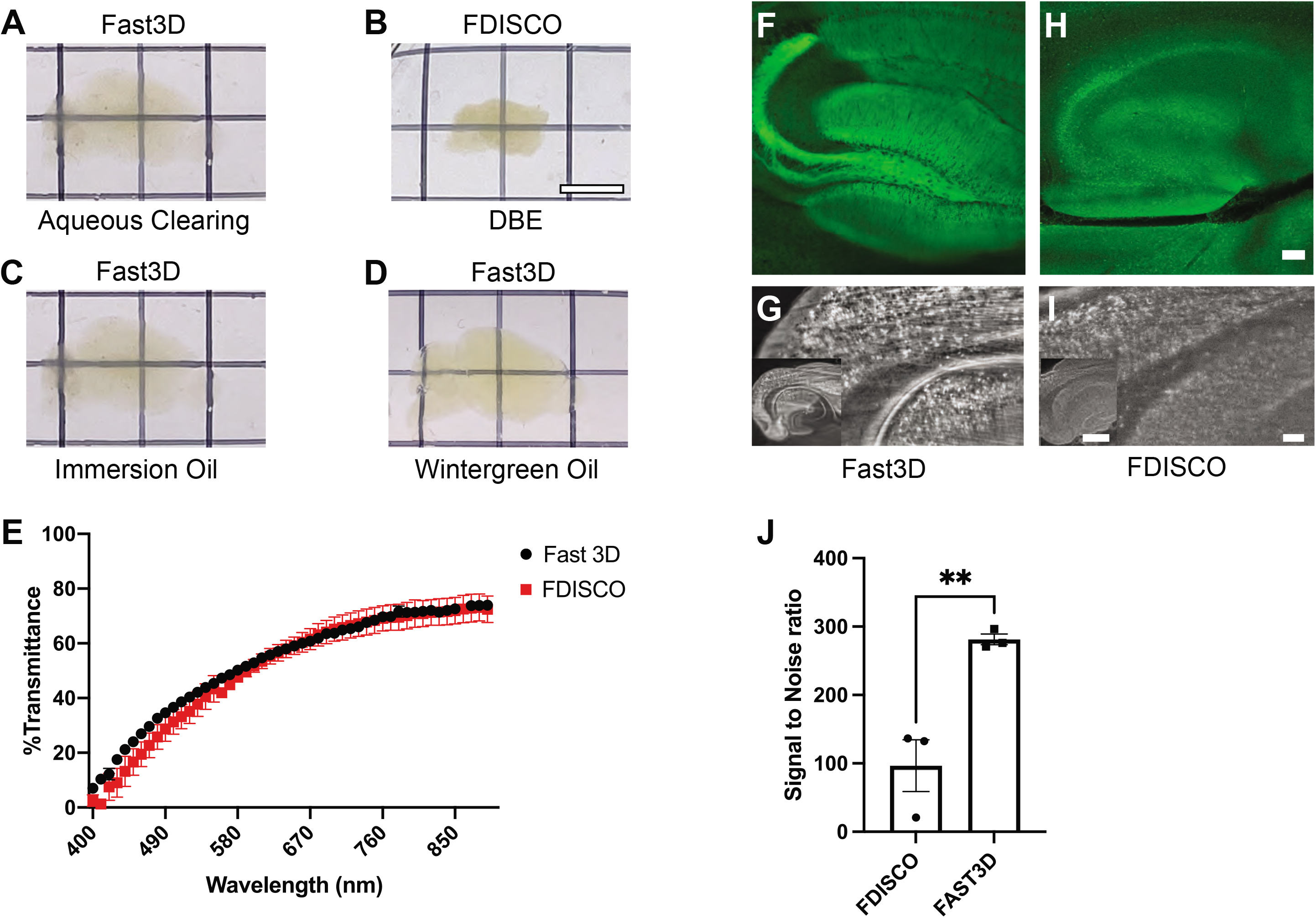
Comparison of Fast 3D Clear with other clearing methods. A-D) Sagittal Images of brain hemispheres subjected to Fast 3D Clear in aqueous clearing solution (A), FDISCO in DBE solution (B), and Fast 3D Clear in immersion oil (C) and Fast 3D Clear in wintergreen oil (D). E) Comparison of light transmittance in brain hemispheres cleared with Fast 3D Clear and FDISCO, 2-way ANOVA F (1, 4) = 1.082, p=0.3571. F-I) Representative confocal and light sheet images from GCaMP3-CaMK2 paired hemispheres cleared with Fast 3D Clear or FDISCO. J) Signal to noise ratio of Fast 3D Clear and FDISCO cleared CaMK2-GCaMP3 hemispheres. Unpaired t-test p=0.0088 (Scale bars 6 mm and 200 μm and 100 μm).

### Fast 3D Clear preserves endogenous transgenic fluorescence in adult mouse brains and whole adult mice and embryos

To test whether Fast 3D Clear can preserve the fluorescence of common fluorescent proteins, we applied our protocol to four transgenic mouse lines: 1) Thy1-GFP-M characterized by very high levels of GFP expression in sparse neuronal populations (Ariel, 2017). 2) GCaMP3-CaMK2-Cre (Tsien et al., 1996; Zariwala et al., 2012), in which GCaMP3 calcium-sensitive fluorescent protein is expressed in CaMK2-positive neurons, 3) tdTomato-VGAT-Cre (Kaneko et al., 2018), in which tdTomato is expressed in inhibitory neurons, and 4) cFos-Cre^ERT2^-tdTomato (Guenthner et al., 2013), in which tamoxifen administration results in tdTomato labeling of neurons active during a behavioral task. In all four transgenic lines, adult brains became completely transparent while maintaining their corresponding fluorescence. Specifically, Thy1-GFP^+^ neurons and GCaMP3^+^ CaMK2 neurons were visualized in the dorsal hippocampus and ventral dentate gyrus (Figure 3A-3D) and in other cortical regions (Supplemental Figure 3A-3C). The labeling intensity of inhibitory VGAT tdTomato^+^ neurons was lower in the dorsal hippocampus and in the dentate gyrus compared to Thy1-GFP and GCaMP3-CaMK2-Cre (Figure 3E, 3F). After fear conditioning, a memory-associated behavioral task, red fluorescence resulting from tamoxifen-induced tdTomato expression was visualized in cFos^+^ hippocampal cell bodies and dendrites (Figure 3G, 3H). When Fast 3D Clear was applied in whole adult GCaMP3-CaMK2-Cre mice, we could visualize GCaMP3 in the mouse retina, olfactory bulb, and spinal cord (Supplemental Figure 3D-3F).

**Figure 3.**
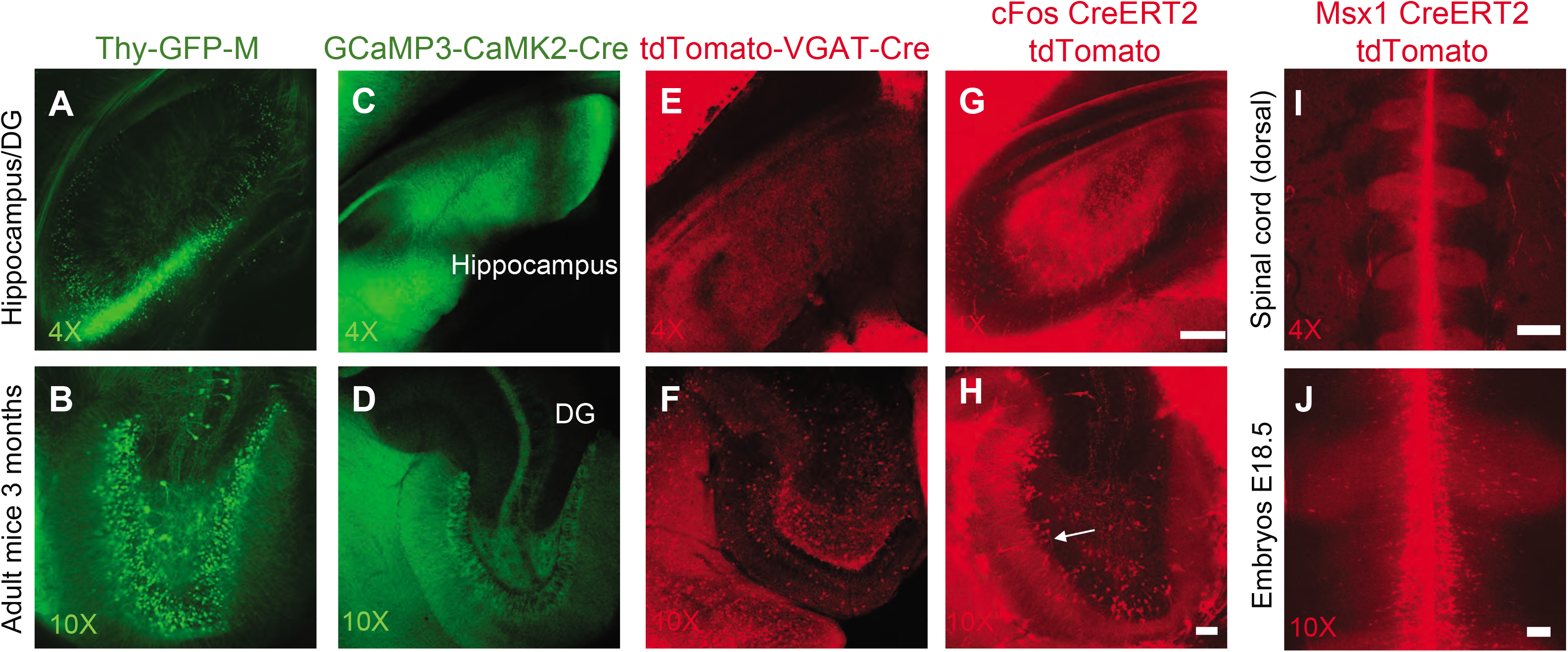
Fast 3D Clear preserves fluorescence in transgenic mouse brains and embryos. A-H) Confocal images taken at 4x magnification (top) and 10x magnification (bottom) showing a dorsal view of the hippocampus of from Thy1-GFP-M (n=3) mice (A-B), GCaMP3-CaMK2-Cre mice (n=4) (C-D), VGAT-Cre-tdTomato mice (n=2) (E and F), cFos-Cre^ERT2^-tdTomato mice after fear conditioning and tamoxifen administration (n=4) (G-H). I-J) Dorsal view of the spinal cord from Msx1-Cre^ERT2^-Tdtomato E18.5 mouse embryos at 4x (I) and 10x(J) magnification (n=4). (Scale bars 200 μm for 4x and 80 μm for 10x magnification).

Similarly, we applied Fast 3D Clear to E18.5 mouse embryos using the Msx1-Cre^ERT2^-tdTomato transgenic mouse line. Confocal microscopy revealed tdTomato labeling in the dorsal and ventral axis of the spinal cord according to Msx1 expression (Duval et al., 2014; Monaghan et al., 1991) (Figure 3I, 3J, and Supplemental Figure 3H, 3I) and in the eye, despite the maintenance of pigmentation (Supplemental Figure 3G). Because some tissues can exhibit high autofluorescence (Croce and Bottiroli, 2014) we visualized the retina (Supplemental Figure 3J-3L) and the spinal cord (Supplemental Figure 3M-3O) from GCaMP3-CaMK2-Cre mice simultaneously at 405, 488, and 568 nm wavelengths (with corresponding emission filters 450/25, 500/30, 585/15) using confocal microscopy. Fast 3D Clear resulted in minimum autofluorescence in the lens of the eye and in the spinal cord, and absence of signal at 405 and 568 nm. These data illustrate that Fast 3D Clear is an optimal approach when maintaining endogenous fluorophores is critical.

### Fast 3D Clear preserves the fluorescence of synthetic and genetically encoded labels at multiple emission wavelengths

To examine whether Fast 3D Clear can preserve a variety of non-endogenous fluorescence of various wavelengths, we delivered fluorescent dyes or virally-expressed fluorescent proteins to the dentate gyrus of the adult mouse hippocampus. Intracranial administration of Fast Blue, a fluorescent dye used as a retrograde neuronal tracer, labeled nuclei in the dentate gyrus and in the entorhinal cortex, a brain region known to provide direct inputs to the dentate gyrus (Witter et al., 2017) (Figure 4A, 4B and Supplemental Figure 4A). Administration of AAV1 CaMK2-GCaMP6 or retro-AAV2 Arch-tdTomato viruses in the dentate gyrus region showed that GFP and tdTomato fluorescent proteins can be preserved and visualized in intact adult mouse brains (Figure 4C-4F and Supplemental Figure 4B, 4C). Next, we created a retrograde virus (retro AAV2) expressing an infrared fluorophore (IRF670). Fast 3D Clear was able to preserve the infrared fluorescence in adult mouse brains (Figure 4G, 4H, and Supplemental Figure 4D). We also measured the signal of each fluorophore from intact adult mouse brains, simultaneously in standard excitation wavelengths (405, 488, 568, 647 nm) with corresponding emission filters (450/25, 500/30, 585/15 and 650 IF) using the same imaging settings for all brains. We found that each fluorophore generated signal restricted almost exclusively to its expected emission band (high SNR), while there was minimal fluorescent signal detected out of band through the other filters (low SNR) (Figure 4I and Supplemental Figure 4E-4H).

**Figure 4.**
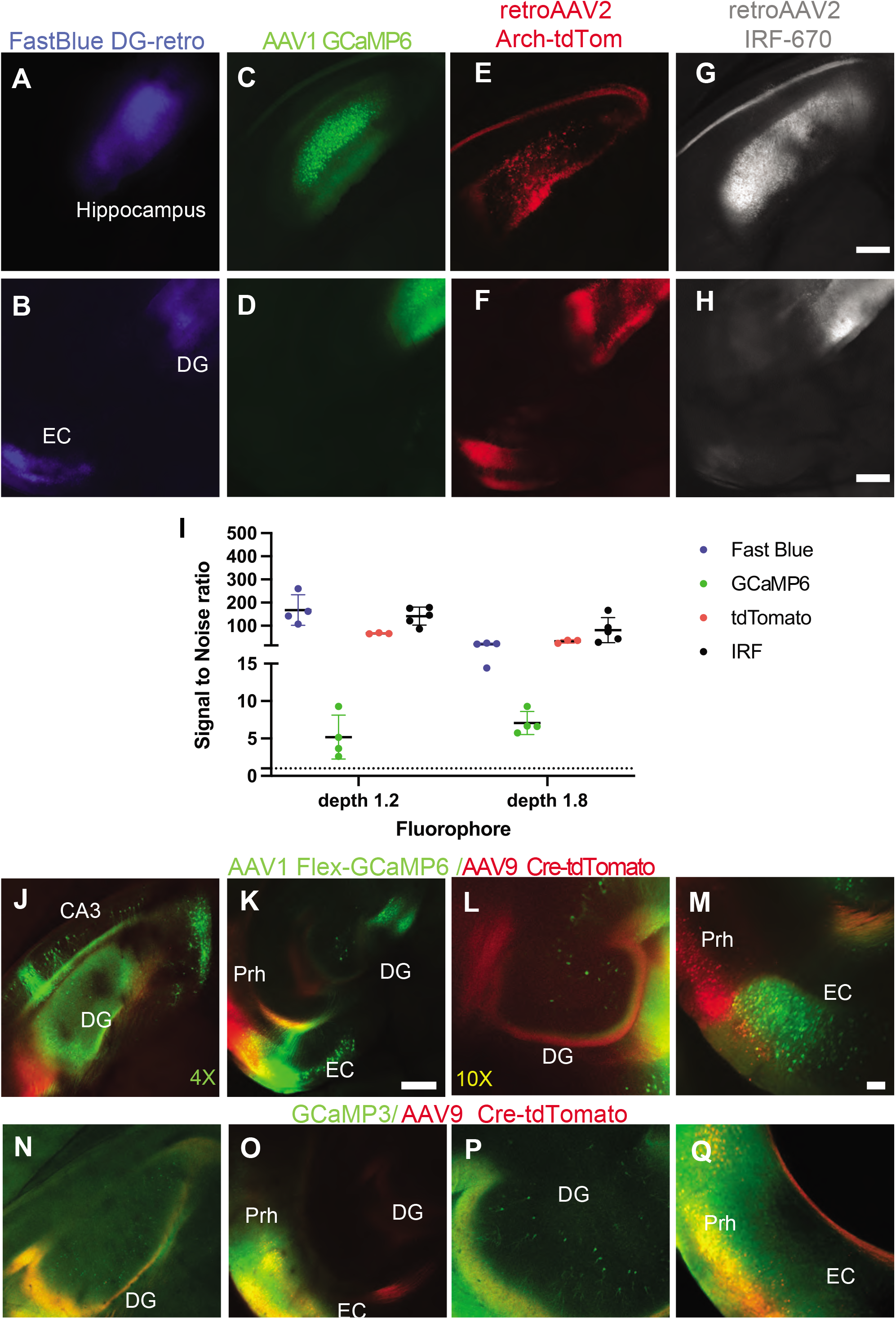
Fast 3D Clear preserves virally-delivered fluorescent proteins and injected dyes. A-H) Fluorescent images of whole adult brains scanned at the same plane displaying the hippocampal formation (top) and the dentate gyrus (DG) and entorhinal cortex (EC) (bottom), from wild-type C57Bl6 animals injected with Fast Blue dye (A and B), AAV1 GCaMP6-CaMK2 (C and D), rAAV2 Arch-tdTomato (E and F) and rAAV2 IRF670 (G and H), (n=7). I) Quantification of signal to noise ratio at 450/25, 500/30, 600/15 and 650LP nm from whole intact mouse brains at two z levels (depth) 1.2 and 1.8mm and representative images. I) 2-way ANOVA for Fluorophores F (3,68)-3504 p<0.0001. J-K) Dorsal hippocampus (J) and Entorhinal/Perirhinal Cortex (K) labeled with GFP and tdTomato fluorophores. L-M) Higher magnifications of the dentate gyrus (L) and entorhinal/perirhinal cortex (M). GCaMP3 homozygous animals injected with tdTomato-Cre virus. N-O) GFP activation in the dorsal hippocampus (N) and Entorhinal/Perirhinal Cortex (O) with cells labeled with both GFP and tdTomato fluorophores. P-Q) Higher magnifications of the Dentate gyrus (P) and Entorhinal/Perirhinal cortex (Q). (N=4). (Scale bars 200 μm for 4x and 80 μm for 10x magnification).

Recently, Zingg et al. discovered that adeno-associated viruses (AAV1/9) exhibit anterograde trans-synaptic labeling properties (Zingg et al., 2017). This application can be used for tracing and visualizing neural circuits in a cell type-and input-specific manner. To expand the application of the Fast 3D Clear method in neurobiology and its utility for studying neural circuits, we injected a Cre-dependent AAV1-GCaMP6 (Ding et al., 2014) virus in the dorsal hippocampus and a tdTomato-Cre-virus in the ventral entorhinal/perirhinal cortex to assess whether an anterograde transfer of AAV virus will activate the Cre dependent expression of the GCaMP6 in the dorsal hippocampus. Three weeks later, in Fast 3D Clear-processed tissue, we observed simultaneous preservation of both GCaMP6 and tdTomato genetically encoded proteins in the same tissue (Figure 4J-4M). Similar results were obtained when we injected the same tdTomato-Cre adenovirus in the ventral entorhinal/perirhinal cortex of GCaMP3 mice (Figure 4N-4Q). We further applied Fast 3D Clear to CaMK2-Cre animals injected in the CA3 hippocampal subregion with a double inverted AAV virus encoding a designer receptor (Gq-mCherry) exclusively activated by designer drugs (DREADD) technology (CNO) (Zhu and Roth, 2014) - a valuable tool to study the function of neuronal circuits, demonstrating the compatibility of Fast 3D Clear with the mCherry fluorophore (Supplemental Figure 4I, 4J). Finally, we found that Fast 3D Clear can preserve fluorescent Cholera Toxin subunit B (CTB). 24 hours after injection into the mouse hippocampus, CTB can be clearly traced into the entorhinal cortex (Supplemental Figure 4K).

Taken together, these data show that Fast 3D Clear is an ideal method to study the structure and function of neuronal circuits. It is an inexpensive, simple, and time-saving method that is compatible with synthetic and genetically encoded fluorescent labels over a broad spectral range.

### Fast 3D Clear is compatible with light sheet and confocal microscopy

We next tested whether Fast 3D Clear is compatible with light sheet and confocal microscopy. Using light sheet microscopy and a Cargille Type A immersion oil-filled cuvette, we visualized cleared brains from the various transgenic lines described above. We show that Thy1-GFP-M mouse brains and CaMK2-GCaMP6-injected brains can be imaged with sufficient resolution at the cellular level (Figure 5A-5C and Supplemental video 1). Additionally, we were able to achieve comparable cellular resolution of the weak GCaMP3 fluorophore to the strongest Thy1-GFP (Figure 5D,5E). Furthermore, we were able to visualize three-dimensionally the cleared brains of cFos-Cre^ERT2^-tdTomato and VGAT-tdTomato transgenic lines, as well as spinal cord from E18.5 mouse Msx1-tdTomato embryos (Figures 5F-5H). Using Imaris software we visualized cleared brains in three dimensions without great compromises to morphology (Supplemental Figures 5A-5C).

**Figure 5.**
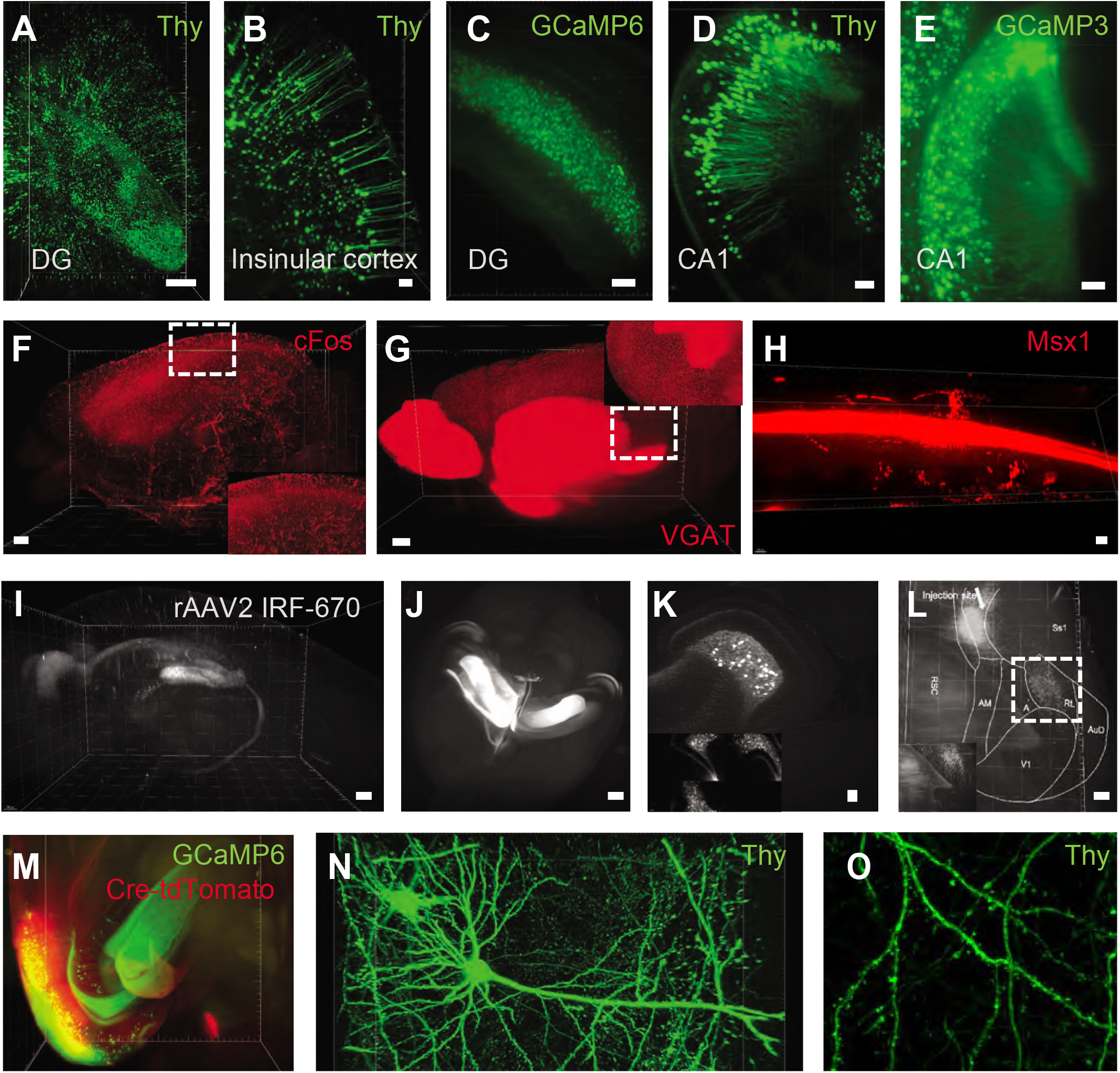
Fast 3D Clear is compatible with light sheet and confocal microscopy. Light sheet images from Thy1 animals in A) hippocampus (Scale bar 1000 μm) and B) cortex (Scale bar 200 μm). C) Light sheet imaging of CaMK2-GCaMP6 injection in the DG (Scale bar 700 μm). D-E) CA1 region of Thy1 animals (D) (Scale bar 100 μm) and GCaMP3-CaMK2 animals (E) (Scale bar 200 μm). F-H) 3D visualization of cleared cFos-Cre^ERT2^-tdTomato brain (F), VGAT-Cre-tdTomato (G) and spinal cord (H) from Msx1-Cre^ERT2^-Tdtomato E18.5 mouse embryos (Scale bar 500 and 200 μm). I) 3D reconstruction of a whole adult mouse brain injected unilaterally with retroAAV2 IRF670 and imaged with light sheet microscopy (2x sagittal view). J) Different brain with retroAAV2 IRF670 (top view) (Scale bar 500 μm). K) Increased magnification (12x) image from the same brain (Scale bar 20 μm). L) Injection of retroAAV2 IRF670 in the S1DZ brain regions showing retro labeled cells (Scale bar 500 μm). M) Light sheet 3D reconstruction from animals injected with tdTomato-Cre (EC/Prh) and GCaMP6 (DG) virus (Scale bar 200 μm). N) 3D reconstruction of cortical neuron from Thy1-GFP animals showing dendritic spines. O) Digital zoom from the same neuron showing spines. (Scale bars N) 20 μm, O) 10 μm.)

We next injected mouse dentate gyri with different adenoviruses, processed the brains with Fast 3D Clear (Figure 5I-5K). In addition to the injected dentate gyrus of the hippocampus, fluorescence was preserved in retrogradely-labeled neurons, and allowed high resolution three-dimensional imaging of structures known to project to the dentate gyrus, including the entorhinal cortex, medial septum, mammillary bodies and projection neurons from the contralateral dentate gyrus (Supplemental videos 2 and 3). We were also able to visualize individual IRF670 hilar cells, with light sheet microscopy (Figure 5K) and confocal microscopy (Supplemental Figure 5D, 5E and Supplemental video 4). Moreover, we were able to detect cFos^+^ in the DG region (Supplemental Figure 5F), Arch-tdTomato (Supplemental Figure 5G) as well as the fine processes of neurons in the CA3 region of the intact mouse hippocampus and retrosplenial cortex after GCaMP6 viral injection (Supplemental Figure 5H, 5I). We also injected the IRF670 virus in the Dysgranular zone (S1DZ) of the somatosensory cortex and visualized cortical areas projecting to S1DZ, such as the rostrolateral area (RL), anterior area (A), anteromedial area (AM) (Figure 5L). Additionally, we could trace in 3D the entorhinal cortex -dentate gyrus neuronal circuit using light sheet (Figure 5M and Supplemental video 5) or a spinning disk confocal microscope (Supplemental Figure 5J). To verify the specificity of the labelling we also processed unlabeled brains of wild-type C57BL6 mice. We found that Fast 3D Clear resulted in minimum background fluorescence (Supplemental Figure 5K-5M).

Finally, in order to examine the ability of our technique to visualize three-dimensionally fine neuronal processes we scanned whole Thy1-GFP-M mouse brains with a 20X lens with high NA (0.95). To this end, we successfully detected spines in two different cortical regions, i.e., insular and retrosplenial cortex (Figure 5N, 5O, Supplemental Figure 5N, 5O and Supplemental video 6).

In summary, we find that Fast 3D Clear is fully compatible with both specialized and simple microscopy instruments, and it provides an efficient solution to three-dimensional imaging of the intact mouse brain.

### Fast 3D Clear is compatible with fluorescent antibody labeling

After applying Fast 3D Clear, whole transparent mouse brains were washed in phosphate buffered saline (PBS) at 4°C for 12 hours. At the end of the wash, all previously cleared specimens returned to their initial opaque state. cFos-Cre^ERT2^-tdTomato-labeled and cleared brain sections (50 μm) were indistinguishable from non-Fast 3D Clear processed brains in terms of both tdTomato fluorescence and overall neuronal structure (Figure 6B, 6E). Lastly, Fast 3D Clear did not hinder the detection of antigens, as demonstrated by the detection of mCherry in the dentate gyrus (Figure 6A-6D), and doublecortin (DCX) in the dentate gyrus, bilateral amygdala, and perirhinal/entorhinal cortex (Figure 6F-6I, and Supplemental Figure 6).

**Figure 6.**
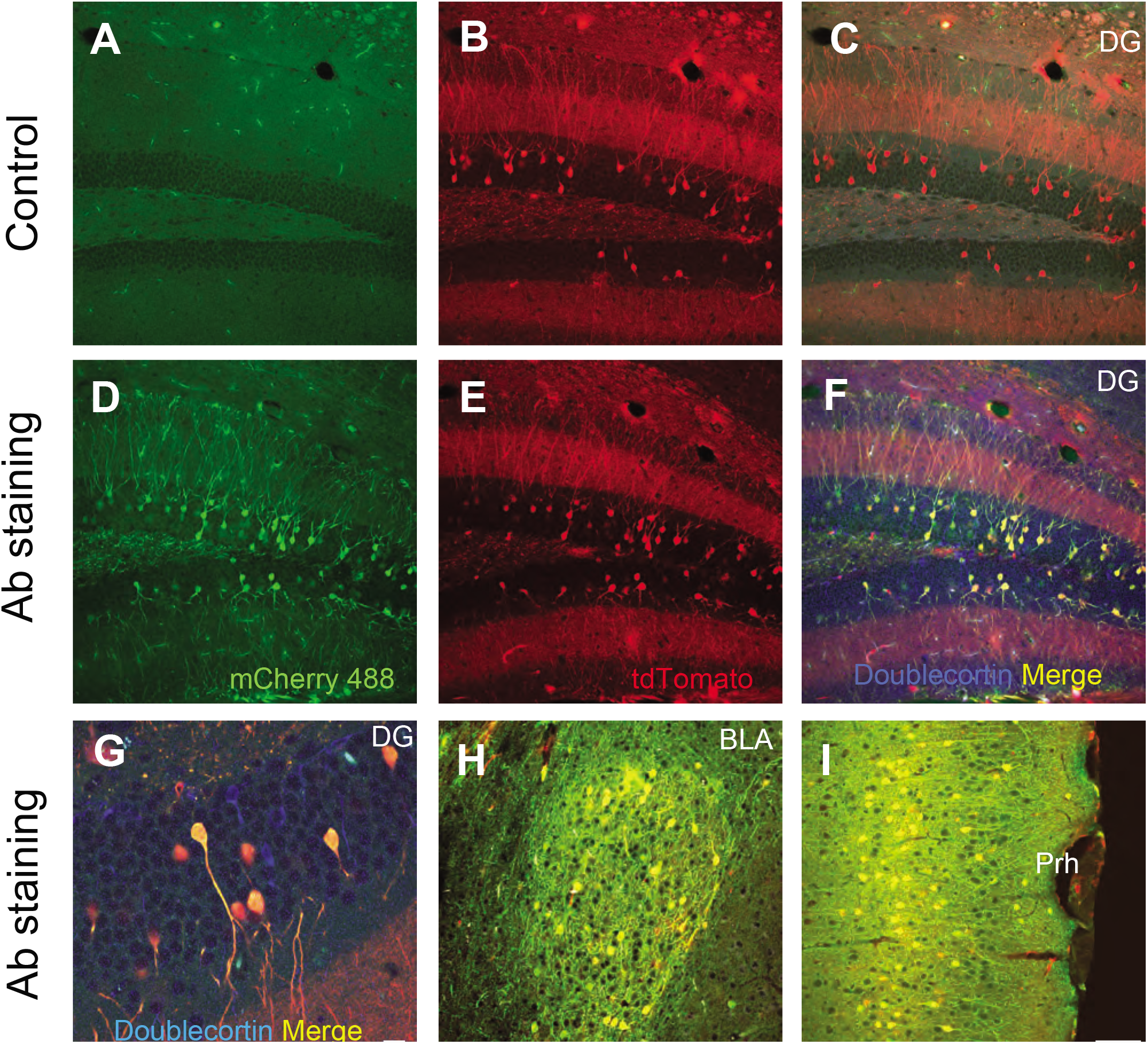
Fast 3D Clear is reversible and compatible with immunohistochemistry. Brain sections after reverse clearing. A-C) Sections from cFos-Cre^ERT2^-tdTomato animals incubated only with secondary antibodies Alexa 488 and 647. Only the endogenous tdTomato (B) can be detected. D-F) Sections from the same animals stained with antibodies against mCherry (488) (D), tdTomato (E) and doublecortin (DCX) (F) (647). G) 60x magnification of the dentate gyrus (DG) stained for mCherry and DCX. H-I) Co-localization of mCherry antibody (500/30) with the endogenous transgenic tdTomato signal in basolateral amygdala (BLA) and in the Perirhinal Cortex, respectively. Doublecortin staining is completely absent from these regions. (Scale bars 20 μm for 20x and 5 μm for 60x magnification).

To test whether Fast 3D Clear is compatible with immunostaining in larger tissue volumes, we cleared brain hemispheres from Thy1-GFP-M mice. After detecting Thy1-GFP^+^cells we reversed the tissue transparency with extended PBS washes and subjected the samples to an iDISCO staining protocol using antibodies against tyrosine hydroxylase (TH) and aGlial fibrillary acidic protein (GFAP) (see methods). Antibody staining was followed by sample incubation in the Fast 3D Clearing solution until full transparency was reached. Fast 3D Clear allowed the preservation of endogenous Thy1-GFP fluorescence, as expected, as well as simultaneous tissue staining for GFAP and TH. Thus, we were able to acquire images from brains labeled with three fluorophores from the dentate gyrus (DG), the central gray of the pons (CGPn), the cerebellum, and the lateral septum (LSI) using confocal microscopy (Figure 7A-7D and Supplemental Figure 7A-L). Additionally, using light sheet imaging or confocal microscopy, we were able to visualize in 3D, triple-stained brain hemispheres (Figure 7E-7H, Supplemental Figure 7M-7U and Supplemental video 7). In a similar fashion, we used GCaMP3-CaMK2 animals and subjected them to a shorter Fast 3D Clear protocol (see Methods). We were again able to preserve the fluorescence in the intact mouse hippocampus (Figure 7I-7K) while maintaining minimum background fluorescence (Supplemental Figure 8A, 8B). The clearing process was reversed, and hippocampi were stained for doublecortin using a short iDISCO protocol (Renier et al., 2016). Although iDISCO caused a small increase in the background fluorescence (Supplemental Figure 8B, 8D), we were able to visualize GCaMP3^+^ neurons and doublecortin^+^ neurons in the dorsal (Figure 7I, 7J and Supplemental Figure 8A-8D and 8G-8L) and ventral dentate gyrus (Figure 7K and Supplemental Figure 8E, 8F), demonstrating that Fast 3D Clear can also be compatible with iDISCO.

**Figure 7.**
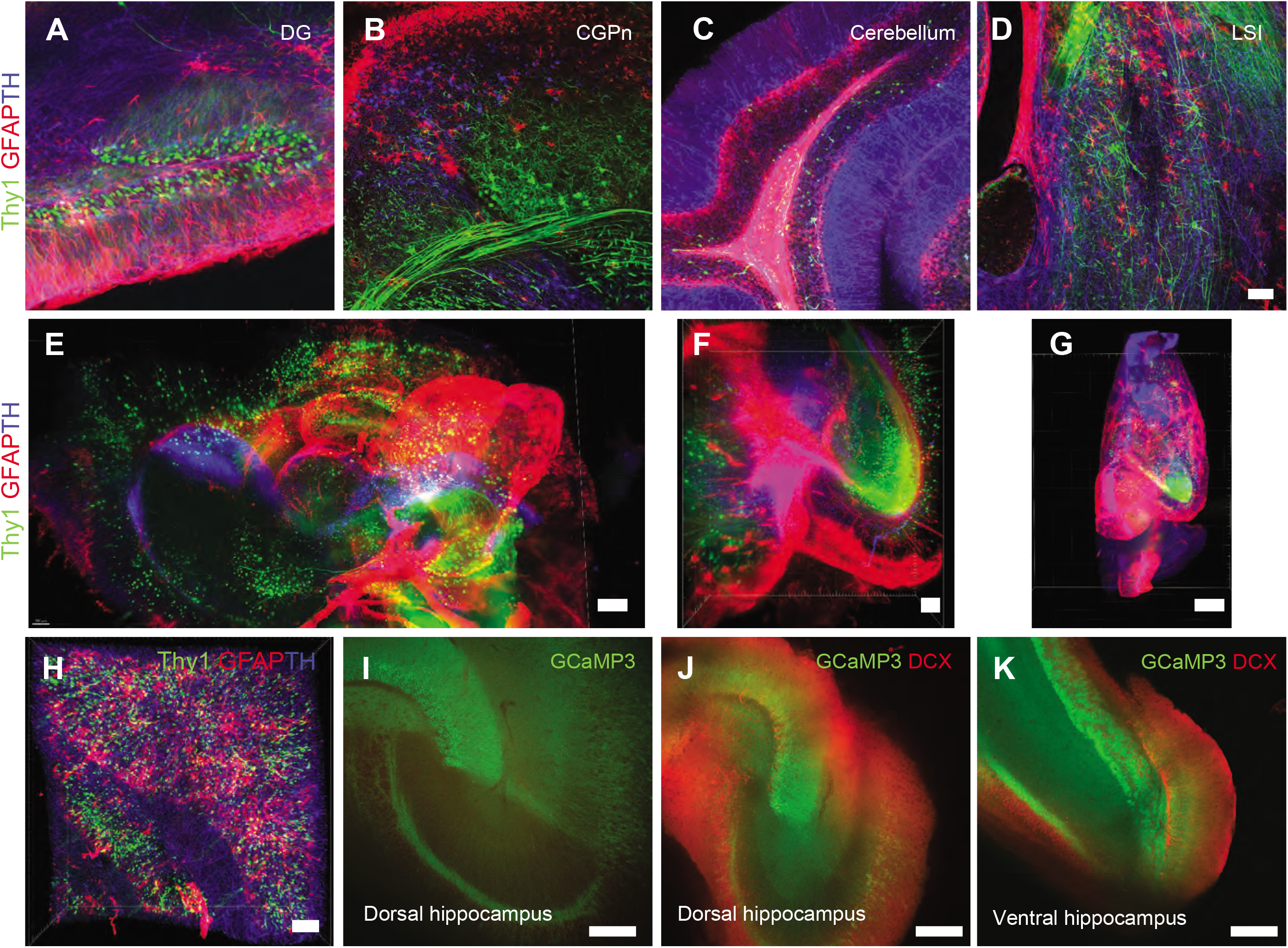
Fast 3D Clear is compatible with whole mount immunohistochemistry. Confocal images from Thy1-GFP-M half brains stained with GFAP (red) and TH (Blue) antibodies A) dentate gyrus (DG), B) central gray of the pons (CGPn), C) cerebellum and D) lateral septum (LSI) using confocal microscopy. (Scale bars 80 μm). E) 3D visualization lLight sheet) from Thy1-GFP-M half brains stained with GFAP (red) and TH (Blue) (Scale bar 500 μm). F) 3D Visualization of a similar brain in larger magnification (5x) (Scale bar 200 μm). G) Top view of another half brain stained with the same antibodies (Scale bar 500 μm). H) 3D visualization of cerebellum from Thy1-GFP-M half brains stained with GFAP (red) and TH (Blue) using confocal microscopy (Scale bar 200 μm). I-K) Whole adult mouse hippocampus from GCaMP3-CaMK2-Cre animals cleared with Fast 3D Clear. I) GCaMP3 fluorescence before staining, J) fluorescence at 500/30 and 650LP nm for GCaMP3 and DCX in dorsal hippocampus and K) ventral hippocampus merged channels (Scale bars 80 μm).

## Discussion

Three-dimensional imaging of cleared tissues has revolutionized neuroscience research, enabling the visualization of single neurons in high resolution and their assembly into circuits in the intact central nervous system. A wide variety of clearing protocols, both organic and aqueous, have become available, and several factors need to be taken into account for choosing the most appropriate approach. These deciding factors include time, cost, complexity, safety, model organism, preservation of fluorophores, and experimental question (Ariel, 2017). Here we describe a new clearing method, Fast 3D Clear, that is compatible with various fluorescent proteins, synthetic dyes, and antibody labeling.

### Advantages of the Fast 3D Clear method

The first benefit of Fast 3D Clear is speed, comparable to sDISCO (Hahn et al., 2019) and FDISCO (Qi et al., 2019). In our hands, intact adult mouse brains, peripheral organs, mouse embryos, and entire young adult mice (P24-P30) become transparent in just three days. Fast 3D Clear is better suited for screening purposes since it is considerably faster compared to other methods, such as CLARITY (Chung and Deisseroth, 2013), iDISCO (Renier et al., 2016), FluoClearBABB (Schwarz et al., 2015), CUBIC (Matsumoto et al., 2019), PEGASOS (Jing et al., 2018), and PACT (Yang et al., 2014).

Second, Fast 3D Clear is inexpensive. Fast 3D Clear does not require specialized equipment, as CLARITY does, and the necessary reagents are only minimal, compared to those required by CLARITY (Chung and Deisseroth, 2013), iDISCO (Renier et al., 2016), FluorClearBABB (Schwarz et al., 2015) and PEGASOS (Jing et al., 2018). In addition, Fast 3D Clear reagents are inexpensive in total and their commercially available quantities are adequate for multiple samples, making our method cost-effective.

Another positive feature of Fast 3D Clear is that it is simple. The protocol includes a limited number of simple incubation steps (50%, 70%, and 90%) with the known dehydration/dilapidation agent THF (THF with BHT inhibitor, Sigma). This particular type of THF, does not require a special process to remove reactive oxygen species (ROS), and is not an immediate hazardous source if handled properly (Hahn et al., 2019; Qi et al., 2019). Fast 3D Clear overcomes the environmental hazard of DBE used in iDISCO (Renier et al., 2016), FDISCO (Qi et al., 2019), and 3DISCO (Erturk et al., 2012), and simplifies the clearing procedure by using an aqueous clearing solution and RI-matching alternatives that do not require complete dehydration with THF. Iohexol has been used previously in RI-matching solutions and as a radiographic contrast agent, and it is an ideal clearing candidate that satisfies our environmental concerns. Moreover, aqueous solutions of high refractive index can be prepared with ease, matching common immersion oils (e.g., Type A, Cargille, RI = 1.515) or reaching close (RI= 1.545) to the RI = 1.562 of DBE at a usable viscosity.

Another limitation of some previously published methods (such as FDISCO and PEGASOS (Jing et al., 2018; Qi et al., 2019)) is tissue shrinkage. Fast 3D Clear completely overcomes the tissue shrinkage. We discovered that the reversal of the dehydration/delipidation THF process can restore tissue size, while extended incubation with water can cause tissue expansion which we can be maintained with the addition of urea without significant fluorescence loss. This moderate expansion feature of our method can be beneficial particularly for the visualization of neuronal circuits.

Using Fast 3D Clear, we were able to maintain the fluorescence of genetically encoded fluorescence proteins for more than a year (GCaMP6, tdTomato, IRF670) and adequately maintain the fluorescence of weaker fluorophores such as GCaMP3. Fast 3D Clear can additionally be used in combination with antibodies to further visualize neuronal circuits. Last, Fast 3D Clear can be used to faithfully reconstruct dendrites from sparsely labeled neuronal structures.

Table 1 provides a summary of comparisons of Fast 3D Clear with similar clearing methods, highlighting its advantages. In short, we believe that Fast 3D Clear is a tissue cleating method that is relatively economical, does not require any special equipment, with low toxicity, fluorescence preservation and immunofluorescence compatibility.

### Limitations of Fast 3D Clear

Despite the strong advantages of Fast 3D Clear over other tissue clearing methods, further testing of some of its applications is still required. First, Fast 3D Clear has not been tested in organisms other than mice, i.e. zebrafish, monkey, and human tissue, to which other methods have demonstrated excellent results (reviewed in (Tian et al., 2021)). For example, although indirect evidence shows that THF treatment is compatible with human embryonic tissue (Belle et al., 2017), further experimentation is needed.

The second limitation of Fast 3D Clear is its modest ability to produce completely transparent hard tissues. While other methods (e.g., BONE clear) (Jing et al., 2018; Wang et al., 2019) are extremely competent, Fast 3D Clear can be applied only to whole young adult mice (P24) with moderate bone clearing. Conversely, application of Fast 3D Clear to 3-month-old mice displays reduced efficiency in hard tissue transparency (bones). It would be extremely useful to test whether we can combine Fast 3D Clear with decalcification and discoloration solutions (as in PEGASOS and BONE clear) to produce a fully transparent adult mouse.

Finally, although Fast 3D Clear is compatible with whole tissue staining (iDISCO) we have not validated the stability of the antibody fluorescence in longer storage conditions (weeks, months) in the Cargille oil or clearing media. We have also not used other immunostaining protocol and reagents, such as antioxidants (i.e., propyl gallate, sDISCO) to evaluate long term storage conditions for antibody treated tissues.

### Outlook

When Fast 3D Clear is paired with light sheet or confocal imaging, it enables the acquisition of high-resolution images to visualize whole brains and embryos. After imaging, the same samples can be processed further with immunohistochemistry with no apparent artifacts compared to non-cleared samples (using the antibodies described herein). Of particular interest is the fact that Fast 3D Clear can be combined with common genetically and virally encoded fluorescent proteins with different promoters (CaMK2, cFos, retro-AAV2), and with the Thy1-GFP-M mouse transgenic line. Our future studies are to further validate Fast 3D Clear by registering the cell types in the Mouse Brain Atlas and, in parallel, perform immunohistochemistry on the same brains in order to bridge the two methods. This will provide an unprecedented strategy for registering multiple cell types both in two and three dimensions, with further applications to translational research. Moreover, it remains of interest to assess whether, besides 3D-imaging and immunohistochemistry, Fast 3D Clear can be combined with other molecular processes such as fluorescent-activated cell sorting and in situ hybridization. In conclusion, Fast 3D Clear can be an important and practical tool with multiple and broad applications in biochemical studies and clinical diagnoses of pathological diseases in the fields of neuroscience, cancer biology and drug screening.

## Author contribution

S.K., A.N. developed clearing protocol; S.K. performed experiments; S.K., A.D., A.L., E.R.K. designed research; S.K., wrote the manuscript with contributions from A.N, A.L., and E.R.K. S.K, AN, A.L. and E.R.K. have filled an IR CU21159.

## Acknowledgements

We thank Dr. Carol A. Mason and Dr. Nefeli Slavi for generously providing us the Msx1 TdTomato embryos. Thy1-GFP-M mouse line was a generous gift from Dr. Josef Gogos. Funding was provided by the Howard Hughes Medical Institute. We would also like to thank the Advanced Instrumentation Core of the Zuckerman Institute and particularly Darcy Peterka, Luke Hammond and Humberto Ibarra Avila for critical comments on the manuscript and technical assistance.

## Methods

### Mice

Mice were maintained under standard conditions approved by the Institutional Animal Care and Use Committee (IACUC) of Columbia University. Animals were housed in a specific pathogen–free animal house under a 12/12-hour light/dark cycle and were provided food and water ad libitum. All wild type mice C57Bl6 were purchased from Jackson Laboratories (8-16 weeks old) (RRID:IMSR_JAX:000664). cFos-Cre^ERT2^ (Luo) (RRID:IMSR_JAX:021882) VGAT-Cre, CaMK2-Cre, GCaMP3 and tdTomato (Ai14) (RRID:IMSR_JAX:007914) were purchased from Jackson Laboratories. Msx1-Cre^ERT2^ mouse embryos were a generous gift from Dr. Carol A. Mason, Columbia University. Thy1-GFP-M mice were a generous gift from Dr. Josef Gogos, Columbia University. Male and female mice were used for all experimental testing. The IACUC of Columbia University Medical Center approved all experiments involving animals.

### Light Transmittance Measurements

The transmittance of half brains cleared with Fast 3D Clear and FDISCO, was measured with a spectrophotometer (Spectronic 200, Thermo Scientific). The brains were placed in a glass cuvette filled with the indicated refractive index matching solution (immersion oil or DBE respectively). Light beam oriented perpendicular to the sagittal plane of the brain and passed through the central part of the brains. Transmittance of light was measured from 400-890nm in 10 nm increments. Light beam size was 7mm. The blank value was measured using a cuvette filled with the indicated refractive index matching solution without sample tissues. Similar measurements were obtained using a second spectrophotometer (Ultrospec 2100 pro) with similar results.

### Signal to noise ratio

To measure signal to noise ratio (SNR) we used GCaMP3-CaMK2 mice brain hemispheres cleared in parallel with Fast 3D Clear and FDISCO methods. Light sheet images were taken at 488 and 568nm from two different planes. Control and experimental areas used, were of the same size. Using ImageJ (Schmidt et al., 2012), we selected two different brain regions that were homogeneously labeled (hippocampus and entorhinal cortex) and measured the fluorescence intensity for 525/50 emission. To calculate SNR we used the formula SNR = (avgForeground - avgBackground) / stdBackground.

### Fluorescence quantification

To measure the fluorescence preservation of Fast 3D Clear we used four fluorophores (Fast Blue, GCaMP, tdTomato, and IRF670.) We calculated the fluorescence intensity in confocal images homogenously labeled in ImageJ using two different depths (z) at 1.2 and 1.8 mm from the top of each whole adult mouse brain (Figure 4). For all the measurements we used the same acquisition settings for all the brains (N=3-5 per condition). All values were normalized with neighboring regions that exhibited near zero fluorescence. The values were expressed as SNR (SNR = (avgForeground - avgBackground) / stdBackground). For long term fluorescence preservation, we used 3-month-old GCaMP3-CaMK2 mice stored in Cargille oil at 4°C for 7 months and freshly perfused mice of the same genotype and age, which were stored in Cargille oil for 7 days (whole brains). We calculated the fluorescence intensity in confocal images, acquired using a 10x 0.4 NA objective type using two different depths (z) at 1.2 and 1.8 mm from the top of each whole adult mouse brain (N=3 per condition).

### Perfusion and tissue preparation

Adult mice were anesthetized with a mixture of ketamine/xylazine (100/20 mg/kg) via intraperitoneal injection. Thereafter, the animals were transcardially perfused with 10 ml phosphate-buffered saline (PBS) and then followed by 30 ml of 4% paraformaldehyde (PFA) in 0.1M NaHPO_4_/NaH_2_PO_4_ or in PBS, pH 7.4. The tissue samples (adult mouse brains) were dissected. For mouse embryo collection, the day of the vaginal plug was defined as embryonic day 0.5 (E0.5), and embryos (E18.5) were removed from anesthetized mothers and transcardially perfused with 4% PFA in PBS. All harvested samples were post fixed overnight at 4°C in 4% PFA and then rinsed 5-6 times with PBS before clearing. For tissue section collection, brains were sliced into 50 μm -thick coronal sections with a vibratome (Leica VT1000 S, Germany).

### Fast 3D Clear protocol

Mice were perfused with ice cold 10 ml PBS and then with 30 ml 0.1M NaHPO_4_/NaH_2_PO_4_ or PBS+4% PFA. In some cases, mice were perfused along with Heparin 10 U/ml in PBS, in order to remove background from blood vessels. Samples were post fixed in 4% PFA overnight at 4°C rotating, end over end. Samples were washed 4-5 times with PBS for 10 min each, followed by 2-3 washes with deionized water at room temperature.

#### Delipidation/Dehydration

Tissues were incubated with THF with BHT (Millipore-Sigma 186562) at 4°C without light exposure, rotating end over end with the following solutions: 50% THF (20ml) in water with 20 μl of triethylamine (pH 9.0) (Millipore-Sigma T0886) for 1 hour, 70% THF (20 ml) in water with 30 μl of triethylamine (pH 9.0) for 1 hour and 90% THF (20 ml) in water with 60 μl of triethylamine (pH 9.0) overnight (∼12-16 hours). pH was measured using pH strips. 20 ml of THF solutions were enough to process 4-5 adult whole mouse brains. At this stage brains shrunk considerably.

#### Rehydration

After overnight incubation with 90% THF the process was reversed rehydrating the samples with: 70% THF (20 ml) in water with 30 μl of triethylamine (pH 9.0) for 1 hour and then with 50% THF (20 ml) in water with 20 μl of triethylamine (pH 9.0) for 1 hour. At this stage, previous shrinkage was reversed.

Finally, samples were washed with ddH20 4-5 times for 10 min each (up to 12h).

At this stage, all specimens became translucent. Specimen size expanded (∼50%) after the water washes.

#### Clearing

Tissues were then transferred in an aqueous clearing solution.

For whole mice (p24) we removed most of the internal organs (spleen, liver intestine) and we used the exact same clearing procedure. Specimens were immersed in the same clearing solution with higher refractive index RI= 1.545, for three days.

For 3-month-old mice we used the same steps as stated above and increased the THF incubation times (50% and 70% THF to 12 hours and 90% THF to 24 hours). A good indication that the whole mouse is adequately processed is the shrinkage of the mouse brain while in the 90% THF.

### Fast 3D Clear solution

The clearing solution consists of 48 g Histodenz (Millipore-Sigma D2158), 0.6 g of Diatrizoic Acid (Millipore-Sigma D9268), 1.0 g of N-Methyl-D-Glucamine (Millipore-Sigma M2004), 0.008 g of Sodium Azide (optional) for 40 ml of solution (0.02% w/v) (Millipore-Sigma S2002) and 10 g of Ultrapure Urea for 50 ml of solution (20% w/v) (Millipore-Sigma 15505). To prepare the solution ∼20 mL distilled water was added in glass bottle with all the chemicals and the powders were dissolved using a stir bar, briefly heating at 37°C to speed up the process. The final volume of the solution is ∼50 ml and the RI 1.512-1.515. Addition of 13 ml of water, instead of 20 ml, produces 40 ml of clearing solution with RI 1.545. Urea can be omitted if the samples are used for other downstream processes, such as antibody staining (as antibodies are sensitive to urea) resulting in a 40ml solution with RI of 1.553. The samples were incubated overnight at 37°C with Fast 3D Clear solution (3-5 ml for a whole adult mouse brain are enough) and rotated end over end. The samples were always submerged completely in the solution. After overnight incubation the samples were completely transparent and stored at 4°C protected from light.

### FDISCO clearing

FDISCO was performed according to (Qi et al., 2019). Briefly, brains were dehydrated with THF solutions (mixed with dH_2_O, pH adjusted to 9.0 with triethylamine) at a series of concentrations 50, 70, 80, and 100 volume % (twice or thrice). Pure DBE was used as a refractive index matching solution to clear tissue after dehydration. All steps were performed at 4°C with slight shaking. During clearing, the tissues were placed in glass chambers covered with aluminum foil in the dark.

### RTF clearing

RTF was performed according to (Yu et al., 2018). Briefly, brains were incubated in RTF-R1, RTF-R2 and RTF-R3 sequentially. Incubation time was 16 hours in each solution. RTF solutions were prepared by mixing triethanolamine, formamide and distilled water by volume. RTF-R1: 30%TEA/40%F/30%W. (30% triethanolamine (TEA), 40% formamide (F), 30% water (W) solution). RTF-R2: 60%TEA/25%F/15%W mixture (60% triethanolamine (TEA), 25% formamide (F), 15% water (W) solution). RTF-R3: 70%TEA/15%F/15%W – (70% triethanolamine (TEA), 15% formamide (F), 15% water (W) solution.

### Imaging

Specimens were imaged with confocal or light sheet microscopy in the clearing solution or they were transferred to Cargille immersion oil type A.

#### Confocal microscopy

Whole brains were submerged in Fast 3D Clear medium or immersion oil. The dorsal hippocampus and dentate gyrus were imaged with an inverted confocal fluorescence microscope (Olympus IX81) equipped with a 4x 0.16 NA Fluar and 10x 0.40 NA Plan-Apochromat objectives. The *z*-step interval was 4.76 μm, pinhole size 80-85μm (auto), PMT settings 600-700 and laser power from 5%-25% depending on the fluorophore. Acquisition settings were 1600X1600 and 10p/sec. We also tested Fast 3D Clear using a W1-SoRa-Yokogawa inverted spinning disk confocal microscope using 10x 0.45 NA Plan-Apochromat objective. For spine visualization we used an XA1R confocal microscope with a 20x 0.95 NA Plan-Apochromat objective. Briefly, 0.5 μm optical sections were acquired in 100μm tissue volume. Laser settings were first tested using unlabeled cleared brains to ensure that no autofluorescence in any wavelength was present. All samples were protected from ambient light. Imaris software was used to produce 3D visualization of cleared samples.

#### Light sheet fluorescence microscopy

Brain samples were imaged using a light sheet fluorescence microscope (UltraMicroscope II Light Sheet Microscope, Miltenyi Biotec, Germany) equipped with an sCMOS camera (Andor Neo), a 2x 0.5 NA Olympus VMPLAPO Plan Apochromat objective lens equipped with a correction lens, and an Olympus MVX10 zoom microscope body. The cleared tissues were mounted on the sample holder and imaged in Cargille immersion oil type A with refractive index R=1.515 (Cargille 16842) in the sample reservoir. Some samples were embedded in 4% low melting agar in the Fast 3D Clear solution and then visualized with confocal or light sheet microscopy. For scanning of whole brains and image acquisition from regions of interest, the *z*-step interval ranged from 4-7μm and the laser power to 5%-25% depending on the fluorophores. Other parameters were: sheet width from 30-100%, dynamic focus from 4-17 depending on the zoom setting and exposure time 100ms. For Imaging tdTomato and IRF fluorophores laser 605/52 and 705/72 were used with laser intensities reaching up to 25% of laser power.

These parameters were tested using unlabeled brains that produced no signal at 30% laser power. Samples in immersion oil (RI∼1.515) or clearing solution can be stored at 4°C protected from light, for at least 6 months (time observed). For 3D visualization, Imaris software was used. None of the figures were processed with algorithms for better visualization, except from the spine images in which a deconvolution algorithm was used.

### Immunohistochemistry

Coronal sections (50 μm) along the entire rostrocaudal extension of the hippocampus were cut on a vibratome and stored in 0.1 M Tris pH 7.4, 30% ethylene glycol, and 30% glycerol at −20°C until further processed. Immunofluorescence was performed on every fifth section. Sections were washed 3 times with 0.1 M Tris pH 7.4, 0.15 M NaCl and 0.3% Triton X-100 (TBSTX) at room temperature. Sections were additionally washed 3 times with TBSTX buffer and blocked for 1 hour at room temperature in blocking buffer (TBSTX supplement with mouse IgG (1 μg/ml) and 1% BSA to reduce non-specific binding of mouse antibodies). Primary antibodies were incubated at 4°C for 24 to 48 hours. After primary antibody incubation, sections were washed at least 3-5 times with TBSTX buffer and incubated with the appropriate Alexa fluorescent secondary antibodies (Invitrogen). Sections were washed 3-5 times with TBSTX buffer and then mounted on Superfrost slides (Fisher Scientific) with Slow fade Diamond mounting solution (Invitrogen) or directly to 3D Clearing solution. Antibodies used: DCX 1:500 Millipore (AB2253), anti-mCherry (Abcam Cat# ab205402, RRID: AB_2722769), Alexa Fluor 488 conjugate (Thermo Fisher Scientific) and Alexa Fluor 647 (Jackson Immunoresearch).

Stereotaxic injections were performed as described previously (Kosmidis, et al., 2018). Mice were anaesthetized with a Ketamine/Xylazine mixture (100 mg/kg) and (20 mg/kg) respectively. Injections were targeted unilaterally to the dentate gyrus (dentate mainly) (−1.96 mm anteroposterior (AP), ± 1.28 mm mediolateral (ML), −1.98 mm dorsoventral (DV)), the entorhinal/perirhinal cortex (−3.15 mm anteroposterior (AP), ± 4.3 mm mediolateral (ML), −3.65 mm dorsoventral (DV)) or S1DZ (−1.0 mm anteroposterior (AP), ± 1.28 mm mediolateral (ML), −0.5 mm dorsoventral (DV)). Injection volumes ranged from 250-500 nl and contained rAAV2, AAV1 viral particles or Fast Blue 0.2 mg/ml (Polysciences 17740). Injection time was 10 min. After the injection the needle remained in place for another 10 min before it was withdrawn.

### Immunolabeling Protocol (iDISCO)

A shorter Fast 3D Clear protocol was applied to hippocampi prior to immunolabeling with the steps described in the iDISCO protocol. Hippocampi from GCaMP3-CaMK2 mice were freshly dissected and fixed in 4% PFA for 20 min in room temperature. Hippocampi were transferred sequentially to 50% and 70% THF solution for 20 min each, and then to 90% THF for 2h at 4°C, Rehydration was achieved after 20 min in 70% and then 50% THF solution. After several washes with water at 4°C protected from light the dissected hippocampi were immersed in Fast 3D Clear solution, at 37°C until cleared (2 hours minimum). Wholemount staining performed according to (Renier et al., 2014) in the same hippocampi. Briefly, reversed cleared GCaMP3-CaMK2 hippocampi incubated in PBS/0.2% Triton X-100/20% DMSO/0.3 M glycine at 37°C for 2 hours, then blocked in PBS/0.2% Triton X-100/10% DMSO/6% Fetal Bovine Serum (FBS) at 37°C for 12 hours. Samples were washed in PBS/0.2% Tween-20 with 10 μg/ml heparin (PTwH) for 1 hour twice, then incubated in primary antibody (DCX-1:250 dilution) in PTwH/5% DMSO/3% FBS at 37°C for 24 hours. Samples were then washed in PTwH 3 times for 15 min, then incubated in Alexa-647 secondary antibody at 1:500 dilution in PTwH/3% FBS at 37°C for 12 hours. Samples were finally washed in PTwH 3 times for 15 min each before clearing and imaging. For half brains we used the same protocol as above. Briefly, we first performed Fast 3D Clearing with Urea and imaged the brains. Then we reversed clearing with PBS and distil water washes for 16 hours at 4°C (4-5 washes). Then we followed the iDISCO staining protocol as above, by extending the incubation times with primary and secondary antibodies for 5 days with constant rotation (20rpm) at 37°C. During the washes (2∼hours) we included a washing step with water to create tissue expansion to presumably aid the antibody penetration. We found that lightly damaged half brains have better and faster antibody penetration than intact ones. The half brains were cleared with Fast 3D Clearing solution without Urea in order to better preserve fluorescent staining. For half brains we used mouse GFAP-Cy3 (Sigma C9205) and rabbit TH (Millipore AB152) antibodies at 1:250 dilution.

### Tamoxifen (TAM)

cFos-Cre^ERT2^-tdTomato animals aged 8–12 weeks were administered 5 mg of tamoxifen (Sigma), suspended in 100 μl 1:1 honey:water mixture by gavage once a day, at least 12 hours apart according to Kosmidis et al 2018 (Kosmidis et al., 2018). cFos-Cre^ERT2^-tdTomato received 2 doses of 5 mg Tamoxifen prior to any other experimental manipulations. For Msx1-Cre induction in mouse embryos, the pregnant mother was injected with 3 doses of 4-hydroxy tamoxifen (2 mg) given at E13.5, E14.5 and E15.5.

### Contextual Fear Conditioning in cFos-Cre^ERT2^-tdTomato mice

Mice were exposed for 4 min to the training context A in which they received 3-foot shocks of 0.7 mA for 2 sec, 1 min apart (Learning). Mice received 2 doses of tamoxifen orally (once every 8 hours) and 24 hours later they were exposed to the same context without any shock (Kosmidis et al., 2018). Five days later mice were processed for Fast 3D Clear.

### HEK293FT cell culture, transfection, and production of rAAV2 and AAV

HEK293FT were purchased from Life Sciences and cultured according to manufacturer’s instructions (R700-07). Viral particles were produced using the HEK293FT cell line according to standard procedures in a BSL-2 safety cabinet according to AAV production protocol. Plasmid vectors that were used: rAAV2-retro helper was a gift from Alla Karpova & David Schaffer (Addgene plasmid # 81070; http://n2t.net/addgene:81070; RRID:Addgene_81070), pAdDeltaF6 was a gift from James M. Wilson (Addgene plasmid # 112867 ; http://n2t.net/addgene:112867 ; RRID:Addgene_112867), pAAV2/9n was a gift from James M. Wilson (Addgene plasmid # 112865 ; http://n2t.net/addgene:112865 ; RRID:Addgene_112865), pAAV-CAG-ArchT-tdTomato was a gift from Edward Boyden (Addgene plasmid # 29778 ; http://n2t.net/addgene:29778 ; RRID:Addgene_29778), pCAG-iRFP720 was a gift from Wilson Wong (Addgene plasmid # 89687 ; http://n2t.net/addgene:89687 ; RRID:Addgene_89687). The viral titers were 10^12^/ml. AAV1-CaMK2-GCaMP6 virus particles were purchased from Inscopix. AAV Gag Flex GcaMP6 (Addgene_100835), AAV hSyn Cre-P2A-tdTomato (Addgene_107738) and pAAV-hSyn-DIO-hM3D(Gq)-mCherry (Plasmid #44361) plasmids were also purchased from Addgene.

### Statistics

Statistics were performed with GraphPad Prism 8 software. Repeated-measures ANOVA (Tukey) and t tests were used as indicated. The results are presented as mean ± SEM. Statistical significance was set at p < 0.05 (*p < 0.05, **p < 0.01, and **p < 0.001)

---

**Supplemental Figure 1.**
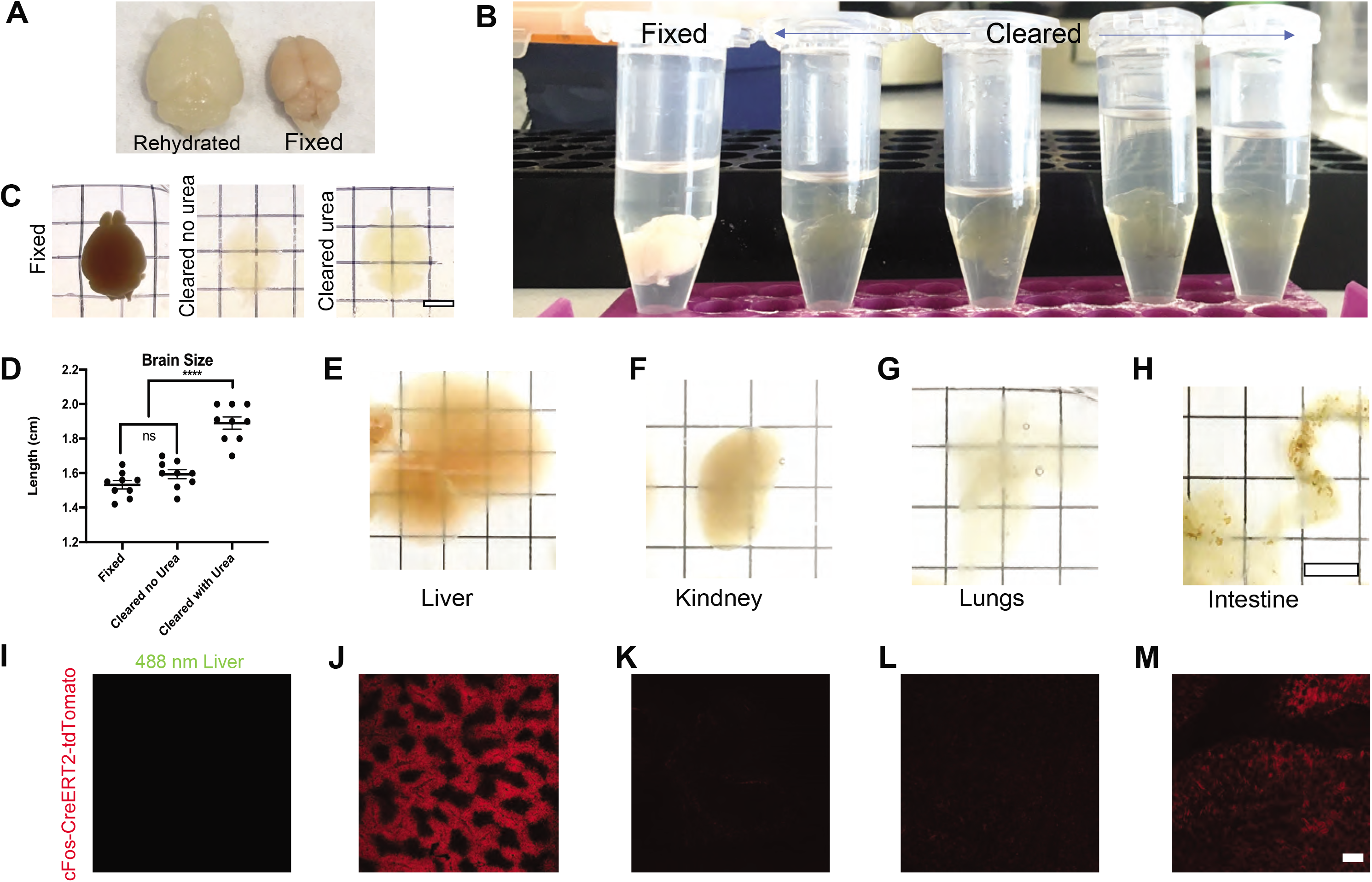
Overview of Fast 3D Clear. A) Images of a rehydrated mouse brain (left) after water immersion and of a fixed brain (right). B) Comparison between fixed and cleared whole brains in Fast 3D Clear solution. C) Representative images of fixed (left), cleared without urea (middle), and with urea (right) depicting size increase. D) Quantification of average mouse brain size (8-16 weeks) after clearing with or without urea compared with fixed brain, One-way ANOVA F (2.24) = 44.11 p<0.0001 between urea and fixed or no urea brains. E-H) Representative images of liver (E), kidney (F), lungs (G), and small intestine (H) cleared with Fast 3D Clear. (Scale bar 6 mm). I) Liver autofluorescence at 500/25 nm from 3-month-old cFos-Cre^ERT2^-tdTomato animals. J-M) TdTomato fluorescence in liver (J), kidney (K), lungs (L) and small intestine (M). (Scale bar 200 μm).

**Supplemental Figure 2.**
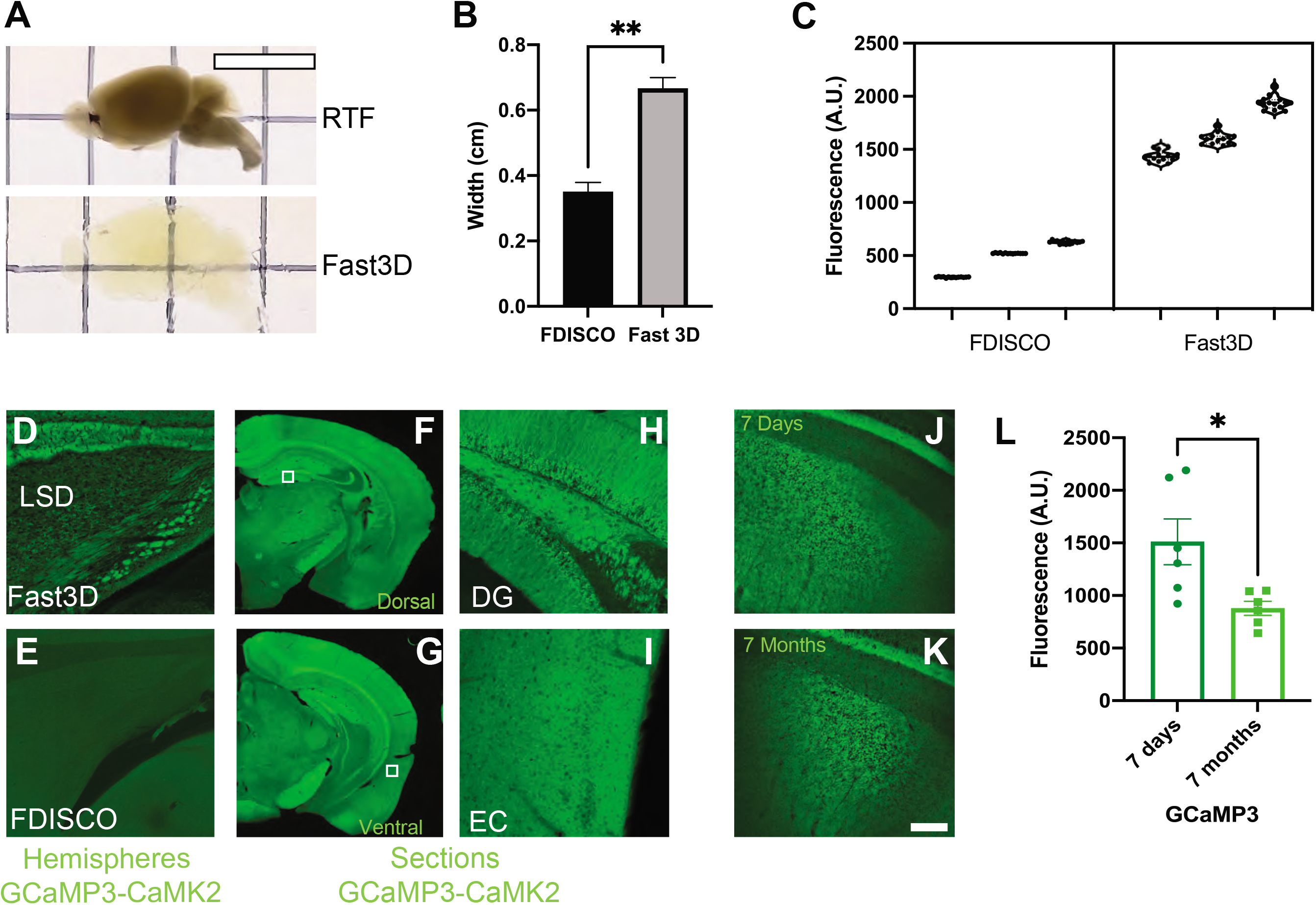
Comparison of Fast 3D Clear with other clearing methods. A) Representative sagittal images of brain hemispheres cleared with RTF (top) or Fast 3D Clear (bottom). B) Quantification of size in cm of brain hemispheres cleared with Fast 3D Clear or FDISCO, unpaired t-test p=0.002, t=7.181, df=4. C) Fluorescence values (A.U) from Figure 2J, nested t-test p=0.0027 two tailed t=6.642, df 4. D-E) Confocal images from lateral septal nuclei (LSD) brain regions of GCaMP3-CaMK2 cleared with Fast 3D Clear and FDISCO. F-I) 50 μm brain sections from GCaMP3-CaMK2 animals showing dorsal (F) and ventral (G) parts of the hippocampus. H-I) Dentate gyrus (DG) and entorhinal cortex (EC) from the same sections. (Scale bar 20 μm). J-K) Confocal mages of GCaMP3-CaMK2 whole brains after 7 days (J) or 7 months (K) in Cargille immersion oil. L) Quantification of fluorescence from 7-day and 7-month GCaMP3-CaMK2 whole stored brains, unpaired t-test p=0.032, t=2.785, df=5.922.

**Supplemental Figure 3.**
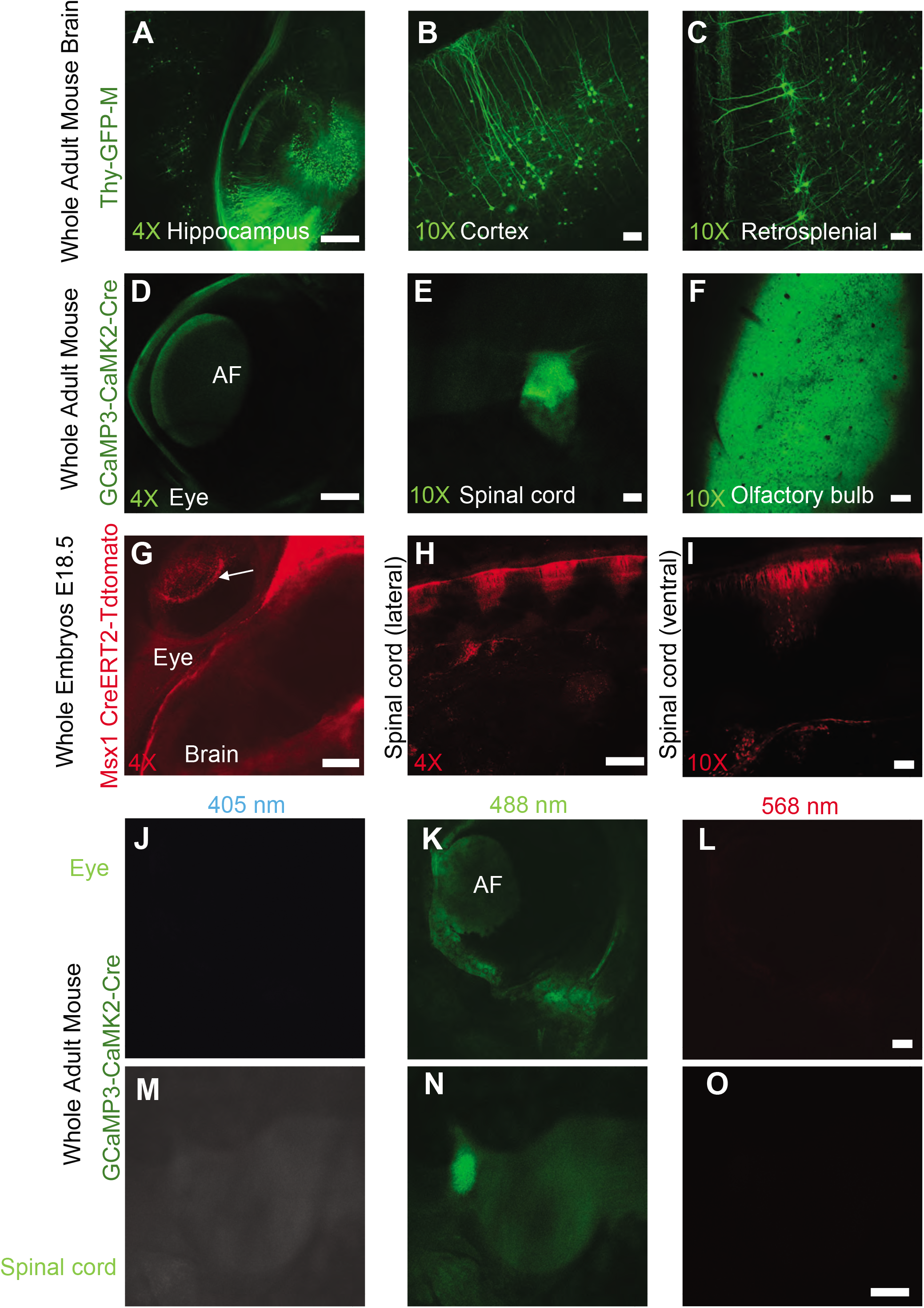
Fast 3D Clear preserves fluorescence in transgenic mouse brains and embryos. A-C) Confocal images from Thy1-GFP-M mouse line showing the hippocampus (4x), and cortical regions at 10x magnification (B-C). D-F) Confocal images from whole GCaMP3-CaMK2-Cre adult mouse, showing eye at 4x magnification (D), olfactory bulb (E) and spinal cord (F) at 10x magnification. G-I) Confocal images from Msx1-Cre^ERT2^-tdTomato E18.5 embryos. In the retina periphery tdTomato+ cells are detected despite the presence of pigmentation (G). Lateral view of the spinal cord at 4x and 10x magnification (H and I), shows tdTomato+ cells and proliferating dorsal progenitors (n=4). J-O) Representative images from the eye (J-L) and from spinal cord (M-O) form adult whole mice showing background autofluorescence (AF) at emission 450/25, 500/30, and 585/15 nm. (Scale bars 200 μm for 4x and 80 μm for 10x magnification).

**Supplemental Figure 4.**
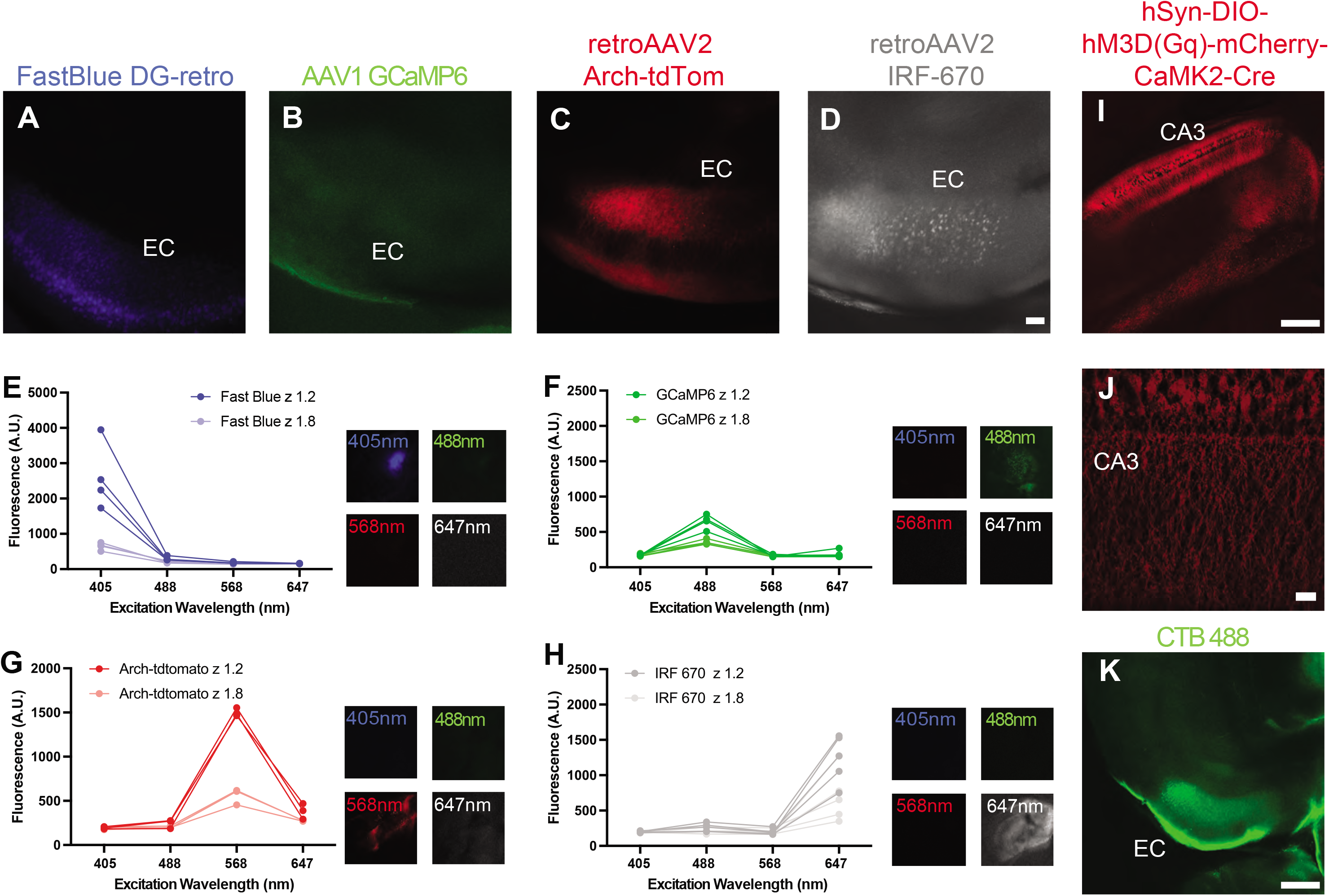
Fast 3D Clear preserves virally-delivered fluorescent proteins and injected dyes. A-D) Fluorescent images at 10x magnification showing labeling in the entorhinal cortex (EC) from animals injected with Fast Blue dye (A), AAV GCaMP6-CaMK2 (control) (B), retroAAV2 Arch-tdTomato (C), and retroAAV2 IRF670 (D). (E-H) Fluorescence (A.U. from Figure 4I). Fast Blue (E), 2-way ANOVA F (3, 18) = 17.24, p<0.0001, GCaMP6 (F), 2-way ANOVA F (3, 18) = 22.94, p<0.0011, tdTomato (G), 2-way ANOVA F (3, 12) = 148.7, p<0.0001, IRF670 (H), 2-way ANOVA F (3, 24) = 5.339, p=0.0058. I) Injection of AAV hSyn-DIO-hM3D(Gq)-mCherry virus (DREADD) in the CA3 hippocampal region of CaMK2-Cre animals and J) higher magnification (N=2). K) Entorhinal cortex region at 4x magnification after injection of CTB in CA1. (Scale bars 200 μm for 4x and 80 μm for 10x magnification).

**Supplemental Figure 5.**
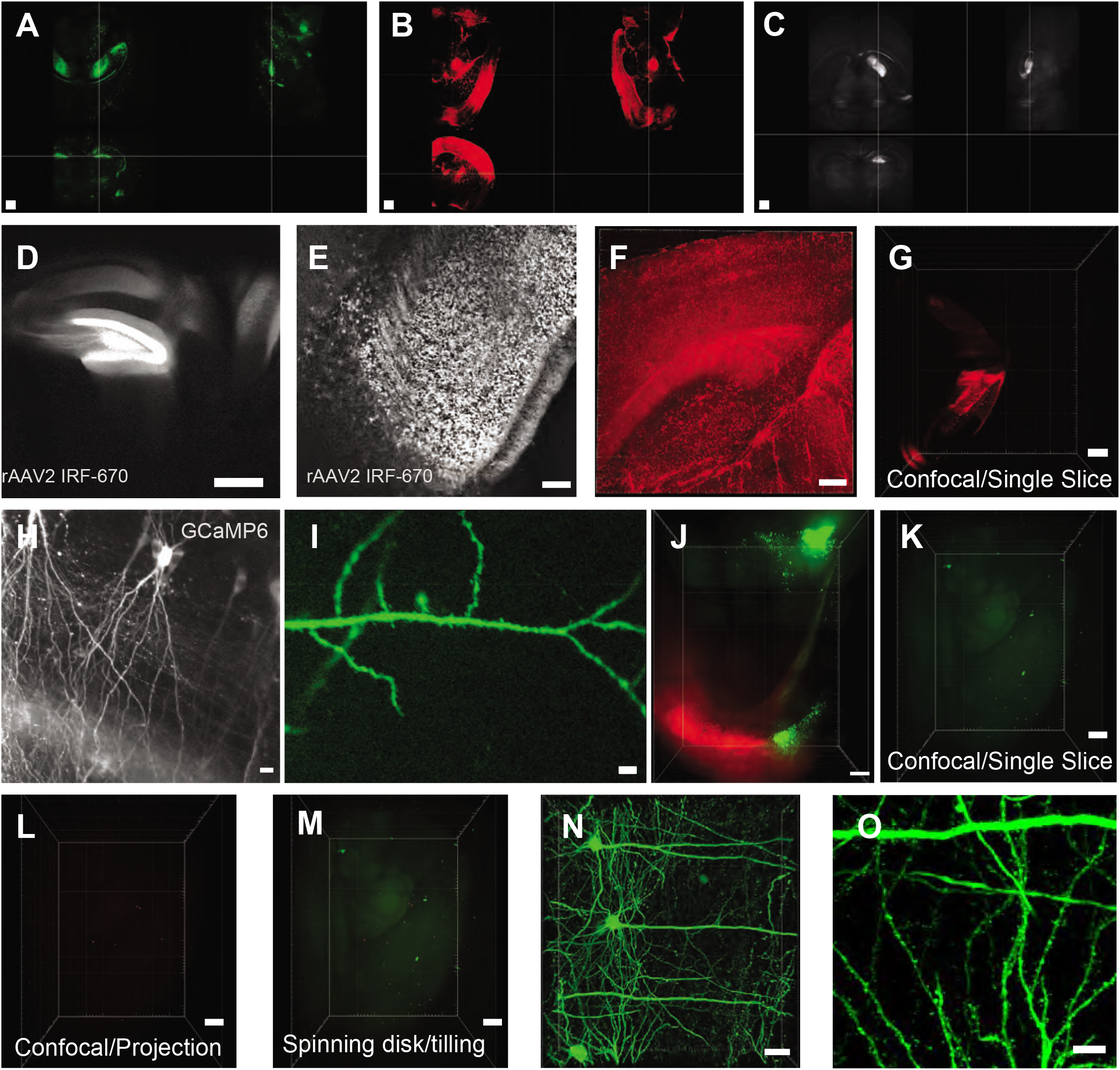
Fast 3D Clear is compatible with light sheet and confocal microscopy. Light sheet images from A) Thy1-GFP, B) cFos-Cre^ERT2^-tdTomato and C) IRF670 in 3 different dimensions (Scale bar 1000 μm). D-E) Single slice from a whole brain injected unilaterally with retroAAV2 IRF670 and imaged with confocal microscopy (4x) (Scale bar 200 μm). (D) Sagittal plane (Scale bar 200 μm), (E) transverse plane (Scale bar 80 μm). F) Maximum projection from cFos-Cre^ERT2^-tdTomato (400mm) showing single cells (Scale bar 200 μm). G) 3D visualization of brain injected with a rAAV2-Arch tdTomato fluorophore (Scale bar 200 μm). H) Single optical maximal projection of sparsely labeled neurons in the CA1 (Scale bar 40 μm) and I) a cortical dendrite labeled with GCaMP6 (Scale bar 20 μm). J) 3D visualization of EC-DG neuronal projection using a spinning disk microscope (Scale bar 200 μm). K-M) Light sheet 3D visualizations from unlabeled C57BL/6 brains showing minimal autofluorescence (Scale bar 700 μm). N) Several cortical neurons from Thy1-GFP-M animals and O) (Scale bar 20 μm) digital zoom region showing spines. (Scale bar 10 μm).

**Supplemental Figure 6.**
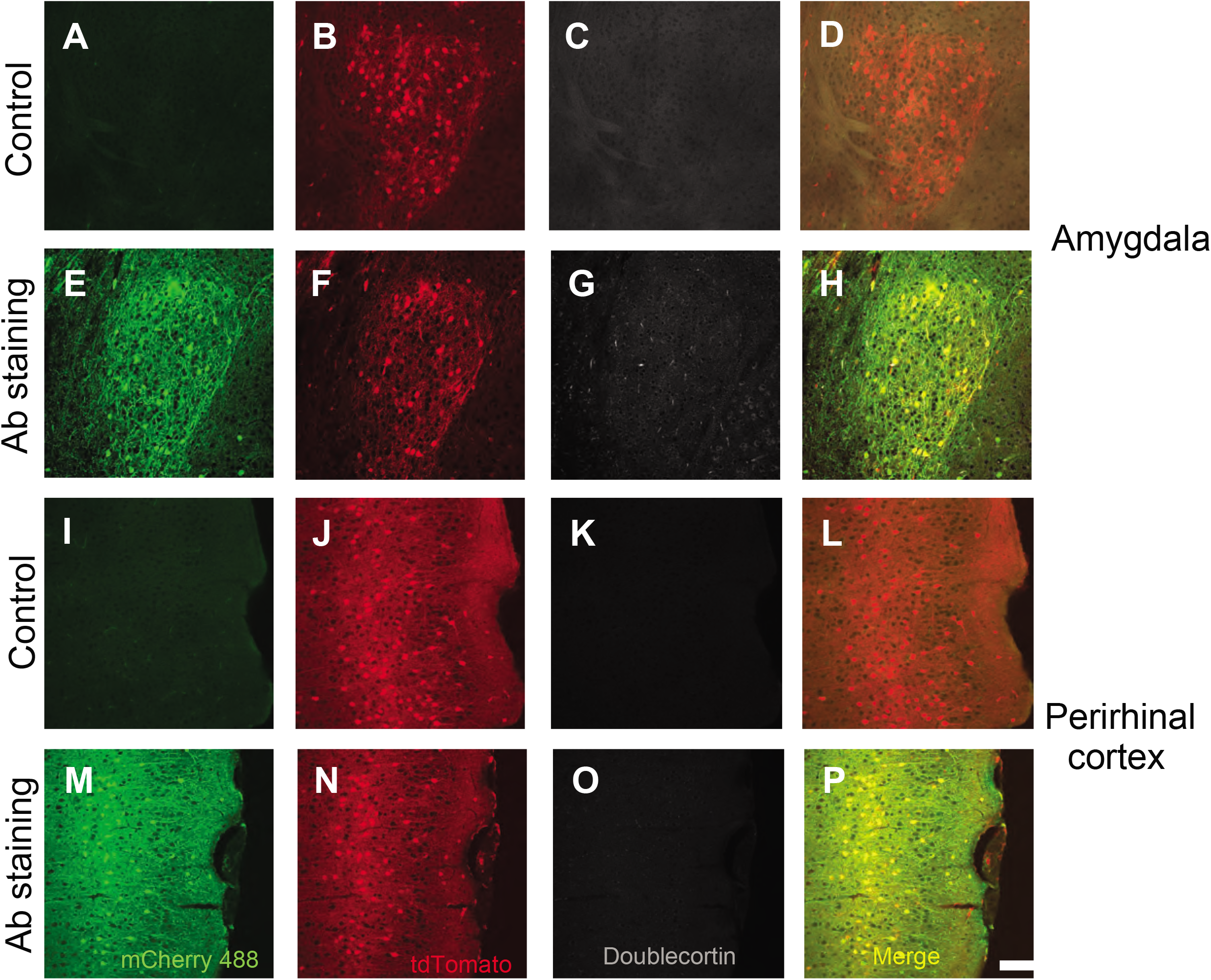
Fast 3D Clear is reversible and compatible with immunohistochemistry. Control (A-D) and (I-L) and stained sections (E-H) and ((M-P) with mCherry and doublecortin DCX antibodies from the basolateral amygdala (BLA) and perirhinal cortex from cFos-Cre^ERT2^-tdTomato animals. (Scale bars 20 μm for 20x magnification).

**Supplemental Figure 7.**
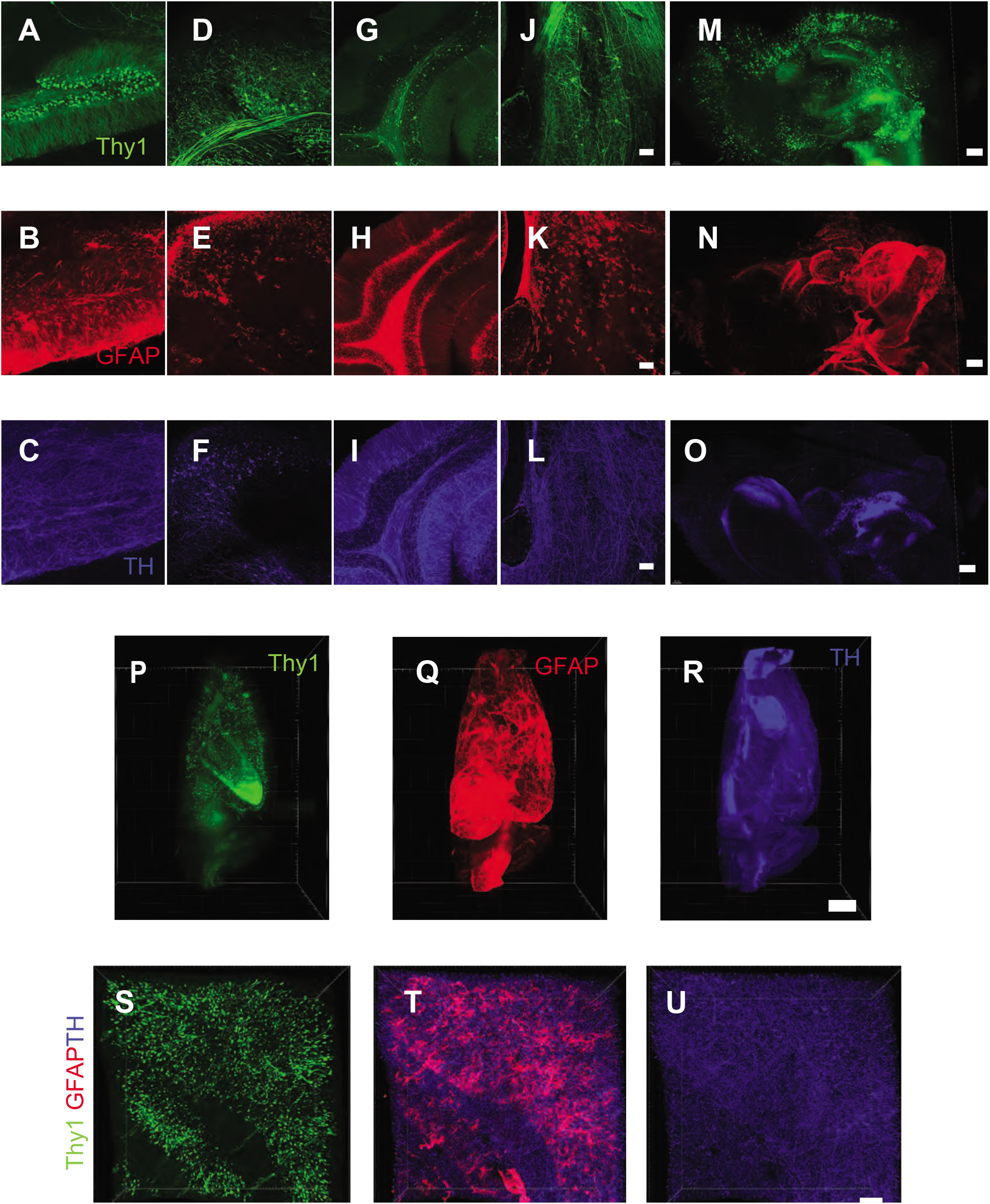
Fast 3D Clear is compatible with whole mount immunohistochemistry. A-U) Single channels from Figures 7A-7H. (Scale bars A-L 80 μm, M-N Scale bars 500 μm, P-Q Scale bar 500 μm, S-U Scale bar 200 μm).

**Supplemental Figure 8.**
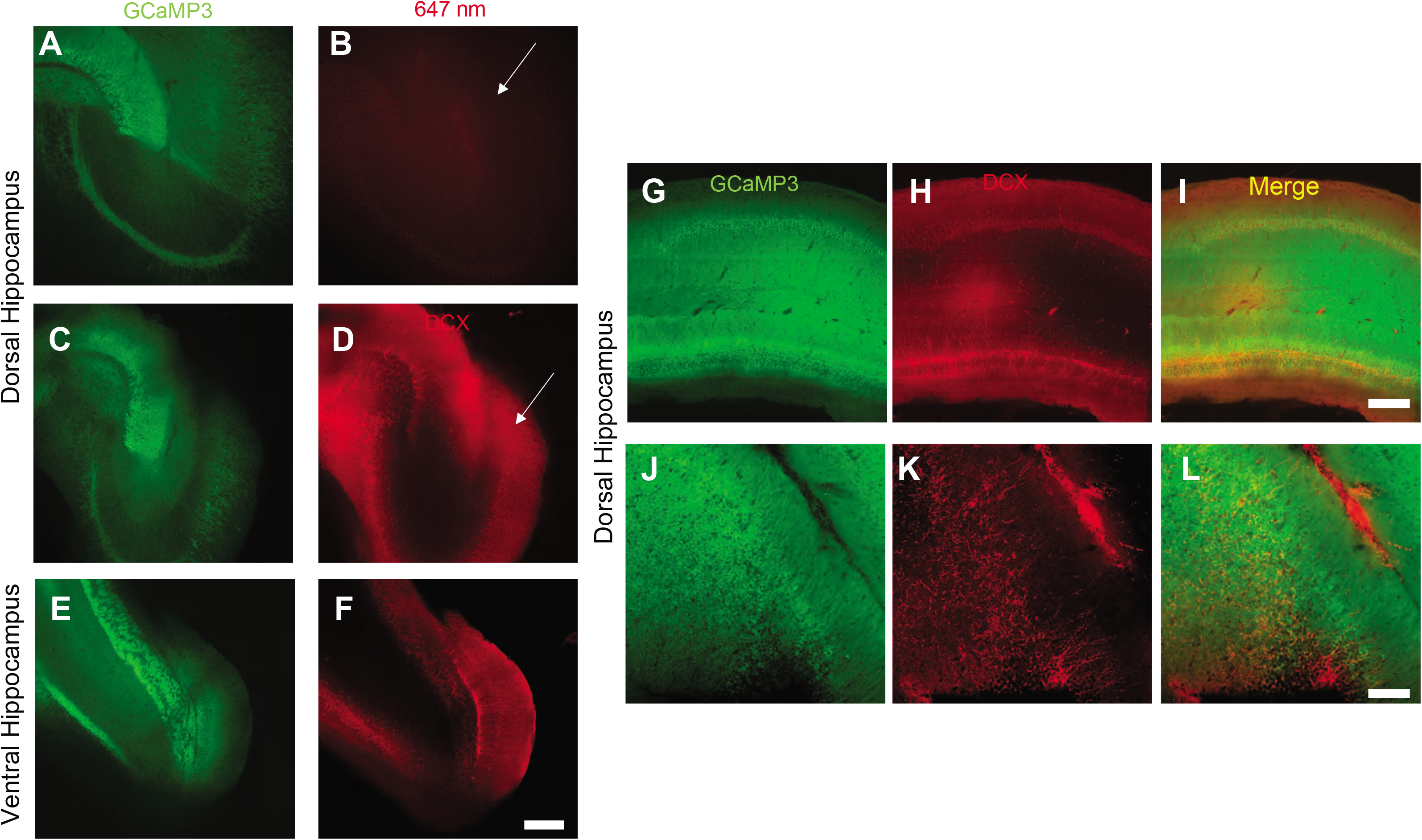
Fast 3D Clear is compatible with whole mount immunohistochemistry. A-F) Single channels from Figures 7I-7K (Scale bar 80 μm). Additional mouse hippocampi after whole mount staining (iDISCO) for doublecortin. G and J) GCaMP3 fluorescence, H) and K) doublecortin (DCX) and I-L) merge plane. (Scale bars 80 μm). Arrows indicate differences in autofluorescence. (N=4).

**Supplemental video 1**. 3D reconstruction/visualization of a Thy1-GFP-M mouse brain.

**Supplemental video 2**. 3D reconstruction/visualization of a whole mouse brain injected with retro-AAV2 IRF670 using light sheet microscopy.

**Supplemental video 3.** Serial sections of a whole mouse brain injected with retro-AAV2 IRF670 using confocal microscopy.

**Supplemental video 4.** 3D reconstruction**/**visualization of 500μm ventral dentate gyrus from a mouse brain injected with retro-AAV2 IRF670 using Light sheet microscopy.

**Supplemental video 5**. 3D reconstruction/visualization of entorhinal cortex projection to the dentate gyrus, using light sheet microscope from mice injected with Cre-tdTomato and GCaMP6 viruses.

**Supplemental video 6.** 3D reconstruction/visualization of a single neuron in the retrosplenial cortex from a whole intact Thy1-GFP-M mouse.

**Supplemental video 7**. 3D reconstruction/visualization of Thy1-GFP-M mouse half brain stained additionally with GFAP and TH antibodies. Note that higher brain regions expressing GFAP protein obscure visualization of regions with lower GFAP^+^ staining.

