## Supplemental Table 1 for "Fast 3D Clear: A Fast, Aqueous, Reversible Three-Day Tissue Clearing Method for Adult and Embryonic Mouse Brain and Whole Body"

| Method | Tissue size | Clearing time | FP preservation | IHC | Tissue size | Toxic | Cell registration | Cost | References |
|---|---|---|---|---|---|---|---|---|---|
| Fast3D | Whole adult mouse brain and whole mouse without skin | 3 days or less (for Juvenile mice also) | Over months | Yes | Normal/ Expansion | + | Not known | ++ | Kosmidis *et al.* (herein) |
| FDISCO | Whole adult mouse brain | 3-4 days | Over months | No | Shrinkage | +++ | No | + | Qi *et al*., 2019 |
| PEGASOS | Whole mouse body with skin | 2 weeks | Over months | No | Shrinkage | ++ | Not known | +++ | Jing *et al*., 2018 |
| iDISCO | Whole-brain immunolabeling | 4 days | Depends on the antibody (idisco.info) | No | Shrinkage | +++ | Yes | +++ | Renier *et al*., 2016 |
